# Phenotypic plasticity reshapes the evolution of species coexistence

**DOI:** 10.64898/2026.11.19.767819

**Authors:** Vishnu Venugopal, Gaurav Baruah, Meike J. Wittmann

## Abstract

Modern coexistence theory (MCT) has been the mainstay of recent research on coexistence of competing species. Competition not only drives evolutionary changes in traits that affect coexistence, but also induces rapid plastic changes. However, a generalised framework that integrates plasticity and evolution into MCT is lacking. We formulate such a framework for a system of two competing species and analyse how breadth of available resources modulates the effect of competition-induced plasticity on the evolution of coexistence components (niche and fitness difference). We find that, in broad environments, divergent plasticity accelerates evolution towards positive invasion growth rates for both species, facilitating coexistence, whereas convergent plasticity hinders this process. Surprisingly, in narrower environments, divergent plasticity can hinder coexistence, leading to exclusion, whereas convergent plasticity can facilitate coexistence. Our results show that plasticity reshapes the eco-evolutionary trajectories to coexistence, highlighting the importance of integrating short-term plastic responses with long-term evolutionary dynamics.

## 1 Introduction

A central goal of ecology is to understand the processes that make species coexist (Cadotte & Tucker, 2017; Levin, 1970; May, 1972). Modern coexistence theory (MCT) (Chesson, 2000a) provides a framework for explaining coexistence through the balance between niche and fitness differences among competing species. Functional traits (and their individual variation) that influence resource acquisition, such as rooting depth in plants (Sankaran et al., 2004) or proboscis length in pollinators (Pyke et al., 2012), determine how differently species (and individuals) exploit resources and thus their potential to coexist. Although trait variation has been extensively studied within the MCT framework (e.g., Barabás & D’Andrea, 2016; Hart et al., 2016; Stump et al., 2022), the role of phenotypic plasticity has largely been overlooked (as also argued by Turcotte & Levine, 2016). Existing eco-evolutionary competition models typically assume traits to be fixed within generations, while studies focusing on the effect of plasticity on coexistence often neglect evolution (Hess et al., 2022). This separation obscures how individual-level plastic responses feed back onto population dynamics and selection, limiting our understanding of coexistence (Hendry, 2016). Here, we develop a generalised eco-evolutionary framework integrating individual-level phenotypic plasticity into coexistence theory to quantify its effect on the evolution of coexistence.

Intraspecific trait variation (ITV) shapes ecological and eco-evolutionary dynamics of competing species (Barabás & D’Andrea, 2016; Begon & Wall, 1987; Gómez et al., 2023; Spaak et al., 2023; van Benthem et al., 2025). ITV arises primarily from genetic variation between individuals and plasticity of individuals in response to environmental heterogeneity (De Bello et al., 2021). Plasticity can rapidly alter ITV (Ashton et al., 2010; Berg & Ellers, 2010; Jackson & Xue, 2024), thereby modifying competitive interactions. Although many studies have examined how plasticity influences ITV under competition (Bennett et al., 2016; Carmona et al., 2019; Novoplansky, 2009; Yang et al., 2024), how such plasticity-mediated changes translate into species coexistence within a MCT framework remains unclear.

Individuals use both abiotic and biotic cues to adjust their traits plastically. In a biotic context, plastic responses depend on interaction partners (Agrawal, 2001; Cahill Jr et al., 2010; Crepy & Casal, 2015; Yoneyama & Bennett, 2024). It can occur reciprocally (Agrawal, 2001; Mougi & Kishida, 2009), only in one partner, be instantaneous or delayed (Dupont et al., 2024; Schneider, 2022; Trappes et al., 2022). For example, some plants exhibit stronger plastic response in specific leaf area and root tissue density to neighbors (biotic cue) than to nutrients (abiotic cue) (Brown et al., 2026). In another example, within-host competition between parasitoids induced plastic shifts in their caste allocation, increasing investment in defence against heterospecific competitors (Harvey et al., 2000). How plasticity in response to biotic cues influences species coexistence has received little attention so far (as argued by Brown et al., 2026; Gibbs et al., 2025; Hess et al., 2022).

Phenotypic plasticity can affect demographic parameters as well as alter competitive interactions by modifying traits associated with resource acquisition, which then can feed back through population densities (Miner et al., 2005). In particular, plasticity induced by competition can substantially alter competitive outcomes (Gibbs et al., 2025; Hess et al., 2022) by driving trait changes in competing individuals (Ashton et al., 2010; Foxx, 2021; Gibbs et al., 2025; Lepik et al., 2005; Lipowsky et al., 2015). These plastic trait shifts are often interpreted as niche differentiation and used to infer coexistence outcomes (Turcotte & Levine, 2016). However, focusing solely on niche differences can be misleading (as argued by Gibbs et al., 2025; Turcotte & Levine, 2016). An accurate method requires accounting for the effects of plasticity on both niche and fitness differences, the determinants of coexistence as per MCT (Chesson, 2000a; Gibbs et al., 2025; Turcotte & Levine, 2016). Two recent works explored empirically the effect of phenotypic plasticity in competing plant species via its effects on niche and fitness differences (Gibbs et al., 2025; Hess et al., 2022). While Hess et al. (2022) found biotic plasticity to promote coexistence, Gibbs et al. (2025) found the opposite. Thus, trait plasticity in response to competition can apparently either promote or hinder coexistence (Stollewerk et al., 2025).

Incorporating evolution by natural selection further complicates how plasticity influences coexistence. Evolutionary responses to competition also shape coexistence through changes in niche and fitness differences (Hart et al., 2019; Pastore et al., 2021). Yet, how competition-induced plasticity interacts with evolutionary change to jointly shape niche and fitness differences remains poorly understood. This lack of clarity partly stems from the still ongoing debate whether plasticity facilitates or hinders adaptive evolution (Ancel, 2000; Chevin et al., 2010; Ghalambor et al., 2007; Grether, 2005; Lande, 2009; Paenke et al., 2007; Pfennig, 2021; Price et al., 2003; Vinton et al., 2022). Moreover, recent empirical work shows that competition-induced plasticity can drive trait convergence or divergence (Bebout et al., 2026). These contrasting plastic responses can have different evolutionary consequences, making the type of plasticity important for predicting its effects on evolution (Ghalambor et al., 2007). Thus, despite growing recognition that both plasticity and evolution influence coexistence, a framework integrating biotic phenotypic plasticity and evolution into coexistence theory is missing.

Here we incorporate competition-induced plasticity in a phenotypic trait associated with resource acquisition into a generalized eco-evolutionary Lotka-Volterra two-species competition model (Barabás & D’Andrea, 2016), explicitly accounting for short-term within-generation trait dynamics. We aim to address the following questions: 1) Does phenotypic plasticity always facilitate the evolution to coexistence? 2) How do resource availability (broad or narrow) and type of plasticity (divergent, i.e., competition-reducing, or convergent, i.e., competition-confronting; Bebout et al., 2026; Gruntman et al., 2017; Novoplansky, 2009) affect the evolution of coexistence?

## 2 Methods

We consider two competing species consisting of individuals varying along a trait axis associated with resource acquisition (figure 1A). Generations are non-overlapping, with each species having a generation time *τ* (see Table 1 for a description of all model parameters). There is a resource availability domain within which the trait of individuals should be for efficient resource utilisation. Individuals, due to competition, adjust their trait via phenotypic plasticity in response to the traits of the interacting individuals and their own trait (i.e., biotic plasticity). Our goal is to find the effect of this plasticity on the eco-evolutionary dynamics of population sizes and the trait of interest. Through this, we aim to understand the effects on the evolution of coexistence via niche overlap and fitness differences. To model this system, we first formulate how plasticity affects the trait dynamics within a generation and then see how the resulting changes in the intrinsic growth rates and competition coefficients alter the evolutionary pressures on the traits of interest and population dynamics.

**Table 1:**
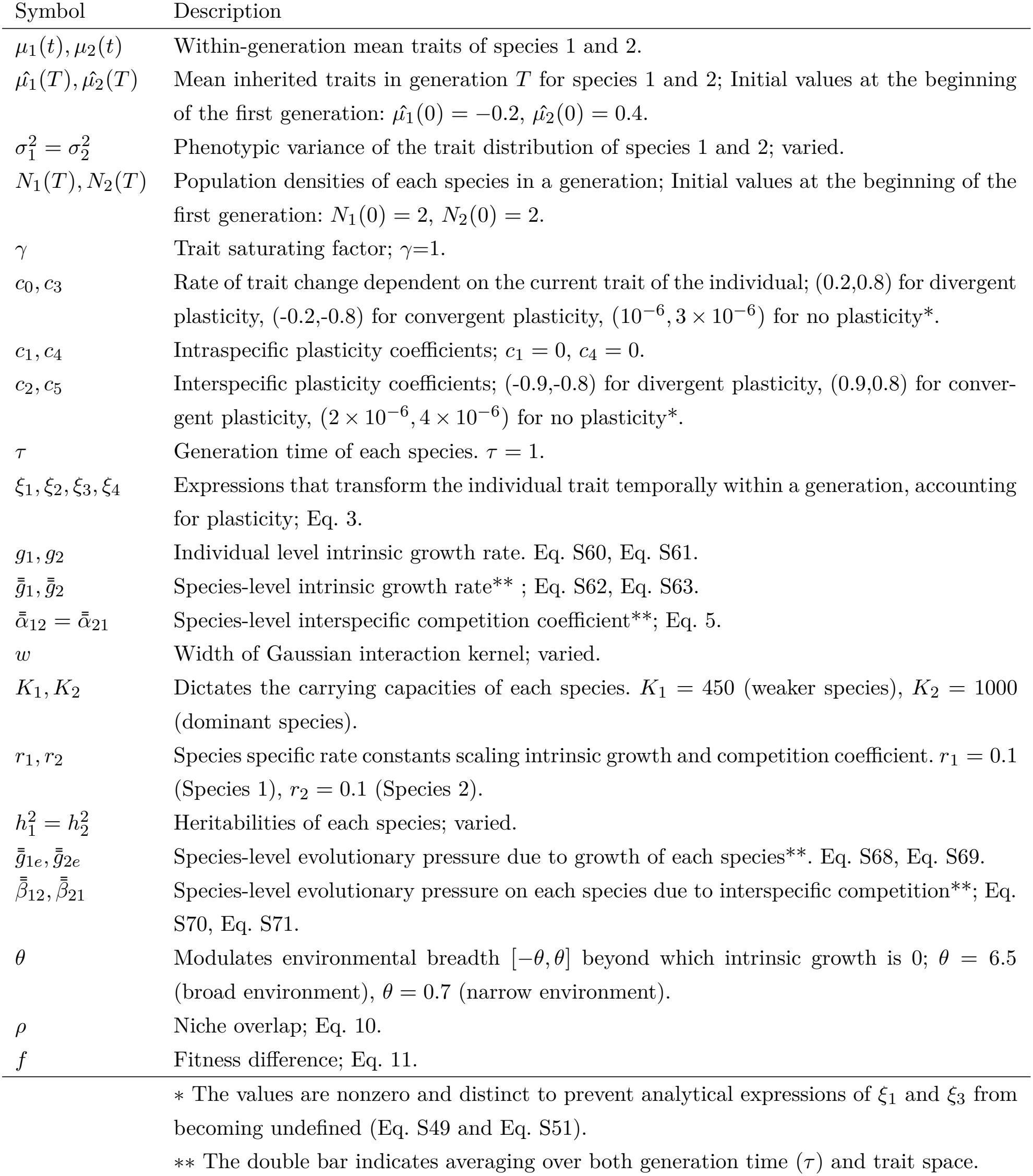
Description of parameters and their values used in the model.

| Symbol | Description |
|---|---|
| *μ*<sub>1</sub>(*t*), *μ*<sub>2</sub>(*t*) | Within-generation mean traits of species 1 and 2. |
| $\hat{\mu_1}(T), \hat{\mu_2}(T)$ | Mean inherited traits in generation *T* for species 1 and 2; Initial values at the beginning of the first generation: $\hat{\mu_1}(0) = -0.2$, $\hat{\mu_2}(0) = 0.4$. |
| *σ*<sub>1</sub><sup>2</sup> = *σ*<sub>2</sub><sup>2</sup> | Phenotypic variance of the trait distribution of species 1 and 2; varied. |
| *N*<sub>1</sub>(*T*), *N*<sub>2</sub>(*T*) | Population densities of each species in a generation; Initial values at the beginning of the first generation: *N*<sub>1</sub>(0) = 2, *N*<sub>2</sub>(0) = 2. |
| *γ* | Trait saturating factor; *γ*=1. |
| *c*<sub>0</sub>, *c*<sub>3</sub> | Rate of trait change dependent on the current trait of the individual; (0.2,0.8) for divergent plasticity, (-0.2,-0.8) for convergent plasticity, (10<sup>−6</sup>, 3 × 10<sup>−6</sup>) for no plasticity*. |
| *c*<sub>1</sub>, *c*<sub>4</sub> | Intraspecific plasticity coefficients; *c*<sub>1</sub> = 0, *c*<sub>4</sub> = 0. |
| *c*<sub>2</sub>, *c*<sub>5</sub> | Interspecific plasticity coefficients; (-0.9,-0.8) for divergent plasticity, (0.9,0.8) for conver-gent plasticity, (2 × 10<sup>−6</sup>, 4 × 10<sup>−6</sup>) for no plasticity*. |
| *τ* | Generation time of each species. *τ* = 1. |
| *ξ*<sub>1</sub>, *ξ*<sub>2</sub>, *ξ*<sub>3</sub>, *ξ*<sub>4</sub> | Expressions that transform the individual trait temporally within a generation, accounting for plasticity; Eq. 3. |
| *g*<sub>1</sub>, *g*<sub>2</sub> | Individual level intrinsic growth rate. Eq. S60, Eq. S61. |
| $\bar{\bar{g}}_1, \bar{\bar{g}}_2$ | Species-level intrinsic growth rate** ; Eq. S62, Eq. S63. |
| $\bar{\bar{\alpha}}_{12} = \bar{\bar{\alpha}}_{21}$ | Species-level interspecific competition coefficient**; Eq. 5. |
| *w* | Width of Gaussian interaction kernel; varied. |
| *K*<sub>1</sub>, *K*<sub>2</sub> | Dictates the carrying capacities of each species. *K*<sub>1</sub> = 450 (weaker species), *K*<sub>2</sub> = 1000 (dominant species). |
| *r*<sub>1</sub>, *r*<sub>2</sub> | Species specific rate constants scaling intrinsic growth and competition coefficient. *r*<sub>1</sub> = 0.1 (Species 1), *r*<sub>2</sub> = 0.1 (Species 2). |
| *h*<sub>1</sub><sup>2</sup> = *h*<sub>2</sub><sup>2</sup> | Heritabilities of each species; varied. |
| $\bar{\bar{g}}_{1e}, \bar{\bar{g}}_{2e}$ | Species-level evolutionary pressure due to growth of each species**. Eq. S68, Eq. S69. |
| $\bar{\bar{\beta}}_{12}, \bar{\bar{\beta}}_{21}$ | Species-level evolutionary pressure on each species due to interspecific competition**; Eq. S70, Eq. S71. |
| *θ* | Modulates environmental breadth [−*θ*, *θ*] beyond which intrinsic growth is 0; *θ* = 6.5 (broad environment), *θ* = 0.7 (narrow environment). |
| *ρ* | Niche overlap; Eq. 10. |
| *f* | Fitness difference; Eq. 11. |

∗ The values are nonzero and distinct to prevent analytical expressions of *ξ*<sub>1</sub> and *ξ*<sub>3</sub> from becoming undefined (Eq. S49 and Eq. S51).
∗∗ The double bar indicates averaging over both generation time (*τ*) and trait space.

**Figure 1:**
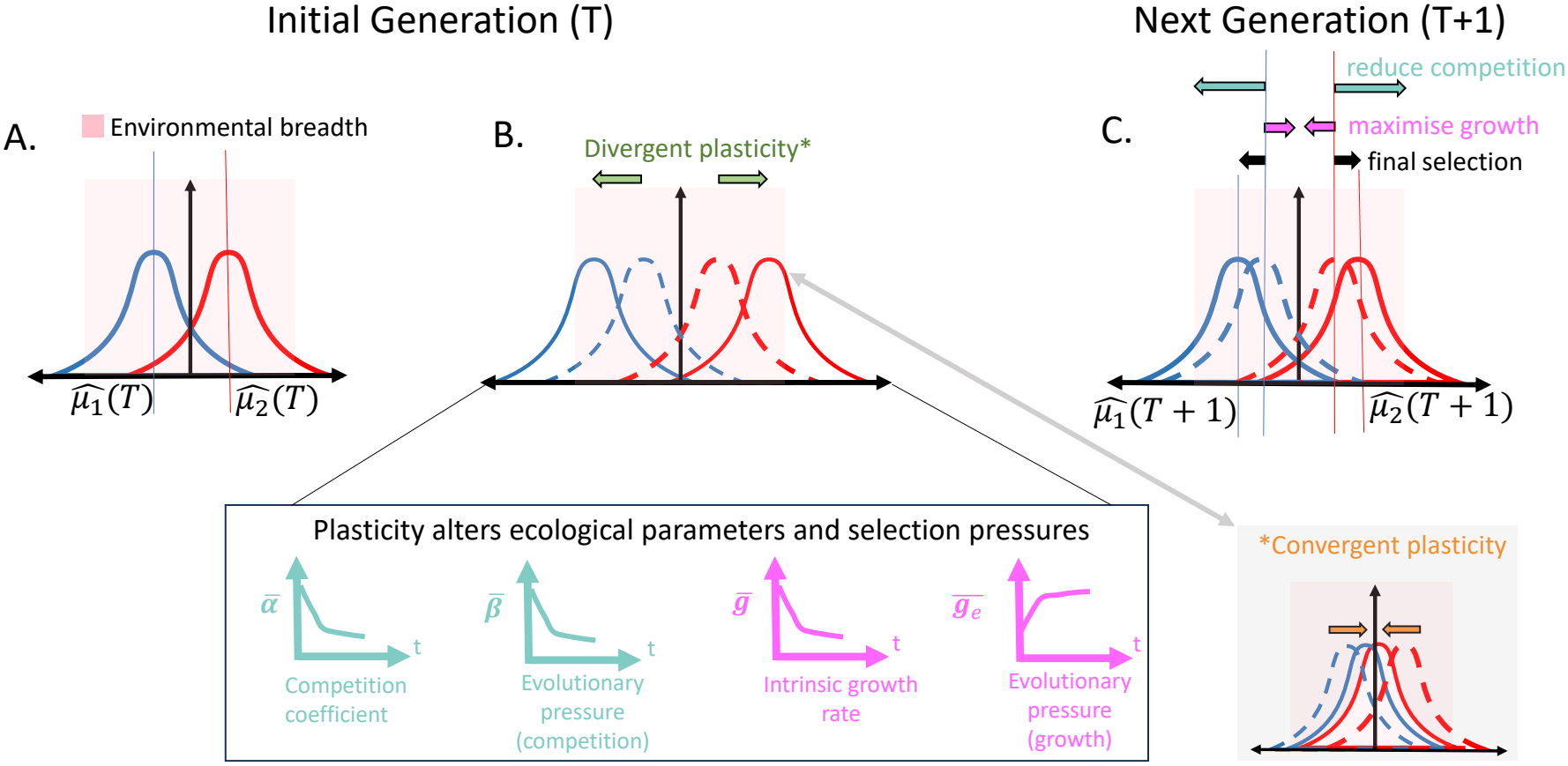
Model overview: Light pink regions indicate the environmental breadth within which individual trait values allow efficient resource utilization. (**A**): Initial phenotypic distribution of two species and mean traits. (**B**): The shift in the distributions of each species due to competition-induced phenotypic plasticity (divergent plasticity is shown here; convergent plasticity is shown as an alternative scenario in the bottom-right). This shift causes temporal changes in the species-level parameters like competition coefficient, intrinsic growth rate, evolutionary pressure to reduce competition and evolutionary pressure to maximize growth, as shown by the four panels under (**B**). (**C**): The species-level parameters in (**B**) are used to calculate the population densities and mean traits of each species after selection for the next generation (see section 2.3 Eq. 6-Eq. 9). Magnitude and direction of the evolution of trait distributions (compared to the original distribution in (**A**)) are shown by the black arrows. This is the sum of the effects of evolutionary pressure on each species to reduce competition (light blue divergent arrows) and to maximize growth (pink convergent arrows), which accumulated over the previous generation.

### 2.1 Trait dynamics due to phenotypic plasticity within a generation

Competition can induce changes in the trait of an individual via phenotypic plasticity. For example, a plant may respond to neighboring roots by growing deeper roots (Gibbs et al., 2025; Guderle et al., 2018; Yang et al., 2024). We denote the phenotypic traits of species 1 and 2 by *a* and *b*, respectively. We assume that these traits are normally distributed, with probability density functions *P*_1_(*a*) and *P*_2_(*b*), means *µ*_1_ and *µ*_2_, and variances 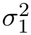 and 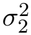, respectively.

Distributions are assumed to be in the quantitative genetic limit (many loci of small effects), under which the distribution remains normal with constant variance under selection (Falconer, 1996). We assume a well-mixed system where all individuals interact. Within a generation, the *i*^*th*^ individual of each species modifies its trait via phenotypic plasticity as given by (see Supplementary section S1.1 Eq. S1 and Eq. S2)

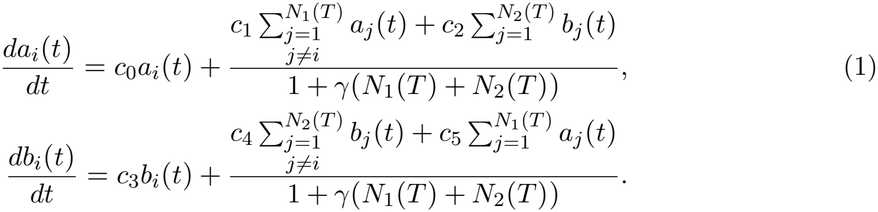

Here, *t* denotes the within-generation time. The rate of change of the trait of each individual depends on its own trait, the traits of individuals of conspecifics and the traits of heterospecifics. *c*_0_ and *c*_3_ are the coefficients that determine the effect of an individual’s own trait in the plastic rate of trait change. *c*_1_ and *c*_4_ are the conspecific plasticity coefficients of species 1 and 2, respectively, and *c*_2_ and *c*_5_ are the heterospecific plasticity coefficients of species 1 and 2, respectively. *N*_1_(*T* ) and *N*_2_(*T* ) are the population densities of species 1 and 2 during the current generation *T* . The plastic change of the trait of each individual is scaled by a term 1 + *γ*(*N*_1_(*T* ) + *N*_2_(*T* )) so that the plastic response saturates with an increase in the population densities and does not go to unrealistically high values in very large populations. Eq. 1 can be rearranged to express the trait change of each individual in relation to the mean traits of each species:

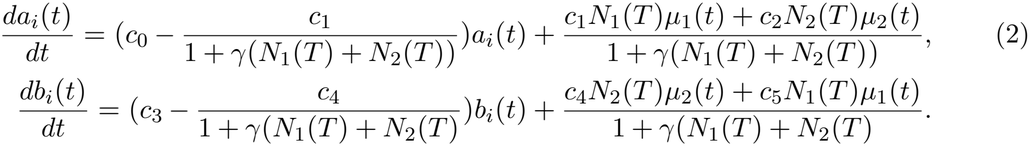

The within-generation mean traits, *µ*_1_(*t*) and *µ*_2_(*t*) in Eq. 2 are defined as 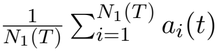 and 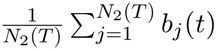, respectively. The mean traits form a system of linearly coupled ordinary differential equations (see Eq. S12, Eq. S13), which are solved analytically to get the expressions for *µ*_1_(*t*) and *µ*_2_(*t*) (Eq. S28, Eq. S29). In the following, we consider only heterospecific plasticity (that is, *c*_1_=0 and *c*_4_=0). Analytically solving Eq. 2, we get the trait of each individual of both species incorporating the plastic modification as given by (see Supplementary Eq. S47, Eq. S48)

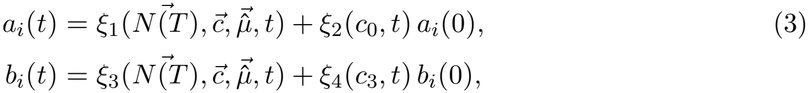

where 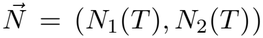 denotes the vector of species abundances in the respective generation, 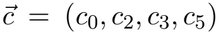 denotes the vector of phenotypic plasticity coefficients, and 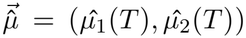 denotes the vector of baseline inherited species mean trait values in the respective generation. Within each generation, plasticity modifies the initial mean traits 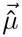 to *µ*_1_(*t*) and *µ*_2_(*t*). The quantities *a*_*i*_(0) and *b*_*i*_(0) represent the initial trait values of the interacting individuals of the species 1 and species 2, respectively, at the beginning of the respective generation. The expressions *ξ*_1_, *ξ*_2_, *ξ*_3_ and *ξ*_4_ are given in the supplementary (Eq. S49 to Eq. S52). These expressions capture the density-dependent dynamics of plasticity. For notational simplicity, we are omitting the index *i* from Eq. 3 hereafter.

### 2.2 Species-level parameters affected by plasticity

Individuals with plastically adjusted traits compete pairwise. At the individual level, we consider a Gaussian interaction kernel to capture the effect of competition. Incorporating the effect of plasticity, the competition coefficient for a trait pair (two individuals) at time *t* is given by

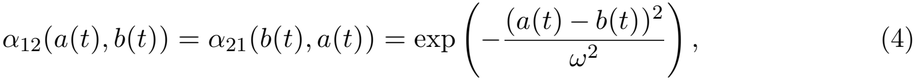

where *ω* is the competition width. At the species level, when each individual adjusts its trait plastically, the competition coefficient can be non-linearly averaged across the generational time and the trait space as follows,

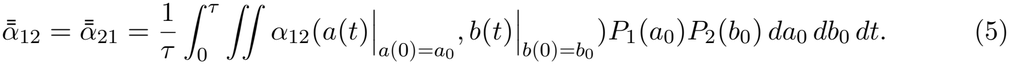

Averaging across the generation time *τ* captures the effect of lifetime plasticity on competition. The intrinsic growth rate of an individual (given by *g*_1_(*a*(*t*)) for species 1 and *g*_2_(*b*(*t*)) for species 2) is a trait-dependent rectangular growth function having a uniform value of 1 when the trait of the individuals falls within the interval [-*θ, θ*], where *θ* is the environment breadth. At the species level, the intrinsic growth rates after accounting for life-long plastic adjustment 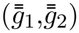 are obtained similarly by integrating individual intrinsic growth rates over the trait space and the generation time (Eq. S62, Eq. S63).

We use a continuous form of the breeder’s equation to quantify evolution (Barabás & D’Andrea, 2016; Pastore et al., 2021). Evolution is quantified as the rate of change of species’ mean trait values driven by the evolutionary pressures acting on each point on the phenotypic axis. Two kinds of evolutionary pressure affect evolution. One is the evolutionary pressure on each individual phenotype to maximize its growth (which at species level takes the form 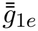 and 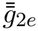 for species 1 and 2, respectively, see Eqs. S68 and S69). The other is the evolutionary pressure on each individual phenotype to reduce competition with individuals of other species (which at the population level takes the form 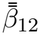 on species 1 by species 2 and 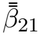 on species 2 by species 1, respectively, see Eq. S70 and Eq. S71). These evolutionary pressures also subsume the effect of within-generation plasticity.

In summary, within a generation, the phenotypic traits of each species are modified as per Eq. 3. This changes all the trait-dependent species-level coefficients, like competition coefficients 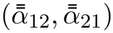, intrinsic growth rates 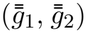 and evolutionary pressures 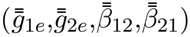, subsuming the effect of plasticity. The species-level coefficients depend on the trait distribution of each species at the beginning of a generation and its plasticity-driven change throughout a generation, the plasticity coefficients, the generation time and the population densities of both species in the generation.

### 2.3 Eco-evolutionary dynamics

The plasticity-embedded competition coefficients 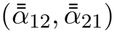, intrinsic growth rates 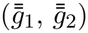 and evolutionary pressures 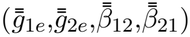 are used in a model for two-species Lotka-Volterra population dynamics and mean trait evolutionary dynamics to approximate the population densities and the mean traits after selection for the next generation. The model takes the form

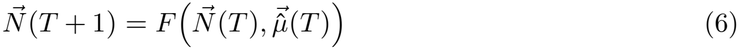

and

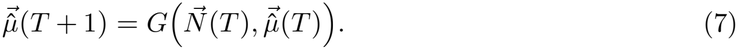

The functions *F* and *G* are obtained as solutions to integrating the following coupled eco-evolutionary system (see Supplementary section S1.2.3) over one generational time interval *t ∈* [0, *τ* ]:

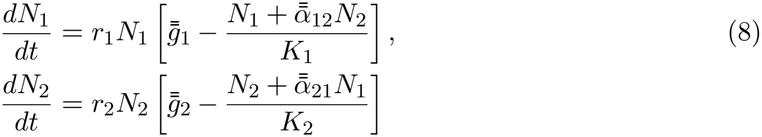

and

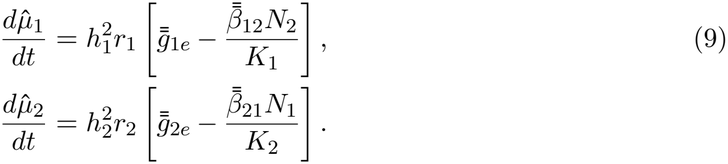

Starting from the initial state 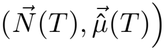, this system is integrated over *t ∈* [0, *τ* ] to get the final state 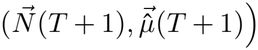. The functions *F* and *G* are then given by the population (Eq. 8) and trait components (Eq. 9) of the solution, respectively. We assume the intraspecific competition coefficients to be 1, assuming that self-limitation is governed by another trait that is independent of the focal trait governing interspecific competition. *K*_1_ and *K*_2_ are the carrying capacities of species 1 and 2, respectively. 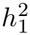 and 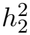 are the heritabilities of species 1 and 2. For each generation *T*, the effect of plasticity (Eq. 3) was recomputed using the population densities 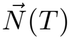 and inherited baseline mean traits 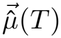 of that generation. The resulting plasticity-embedded species-level coefficients were then used in the eco-evolutionary equations to obtain 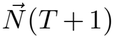 and 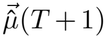. This procedure is iterated across 1000 generations. We use a continuous-time Lotka–Volterra model for population dynamics (Eq. 8) and the breeder’s equation (Eq. 9) for trait evolution, as this framework allows coexistence to be analysed using the well-established niche and fitness difference formulations (Barabás & D’Andrea, 2016; Hart et al., 2019).

### 2.4 Niche difference and fitness difference

Niche difference is the ratio of the strength of intraspecific to interspecific competition. This determines the degree to which a species benefits when being reduced to very low densities. For the population dynamics equations we used, the niche difference (with *ρ* being the niche overlap) is given by (see Supplementary section S1.4 for details)(Chesson, 2018; Pastore et al., 2021)

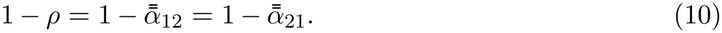

The fitness difference

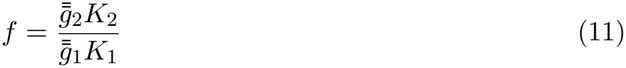

gives the ratio of fitness of species 2 over species 1. An increased fitness difference results in the competitive exclusion of the species with lower fitness if not balanced by the niche difference between species.

### 2.5 Invasion growth rates

The invasion growth rate of a species is the per-capita growth rate of a species when its population is at low density, while the competitor population is at equilibrium. For the population dynamics equations we used, the invasion growth rate of each species is given by (see Supplementary section S1.3 for details)

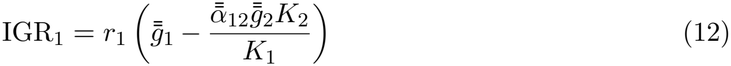

and

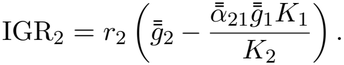

### 2.6 Evolution of coexistence under environmental constraints and plasticity scenarios

The coexistence region is defined by the region obeying the inequality 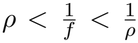 (Chesson, 2000b). For each generation, the niche difference, fitness difference and invasion growth rates are computed as per Eq. (10), Eq. (11) and Eq. (12), respectively. The interaction coefficients used to calculate these are nonlinearly averaged temporally over the generation time *τ* and over the trait space of the phenotypic distribution of each species. We examine how plasticity and environmental breadth influence the evolution of these coexistence metrics. We consider three scenarios for plasticity: no plasticity, divergent plasticity (competition-reducing), and convergent plasticity (competition-confronting) (as observed empirically in Bebout et al., 2026; Gruntman et al., 2017; Novoplansky, 2009). We define divergent (competition-reducing) plasticity as a plastic response that causes the trait distributions to diverge and decreases the species niche overlap, while convergent (competition-confronting) plasticity was a plastic response that increases the niche overlap. These three plasticity scenarios are controlled by the plasticity coefficients, 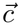 (see Table 1). Regarding environmental breadth, we consider broad environmental breadth to be one that incurs negligible fitness loss on individuals upon trait divergence (*θ >> σ*). Narrow environmental breadth is one where fitness penalties emerge at smaller trait deviations (*θ ≈ σ*). Heritability values of *h*^2^=0 and *h*^2^=0.5 are used to study the cases of whether evolution was absent or present, respectively. A previous study (Pastore et al., 2021) on the evolution of niche overlap and fitness differences have found that environmental breadth (*θ*) and competition width (*ω*) strongly influence the magnitude and direction of niche overlap and fitness differences.

## 3 Results

### 3.1 Role of plasticity in a broader environment

In broad environments (*θ* = 6.5, *ω* = 1.1), divergent plasticity promoted evolution toward coexistence by increasing niche differences, whereas convergent plasticity hindered this process (figure 2). Evolution was mainly driven by evolutionary pressure to reduce competition (solid lines in figure S1A-F) (i.e. 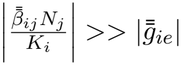 with 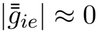 for species *i* in Eq. 9).

**Figure 2:**
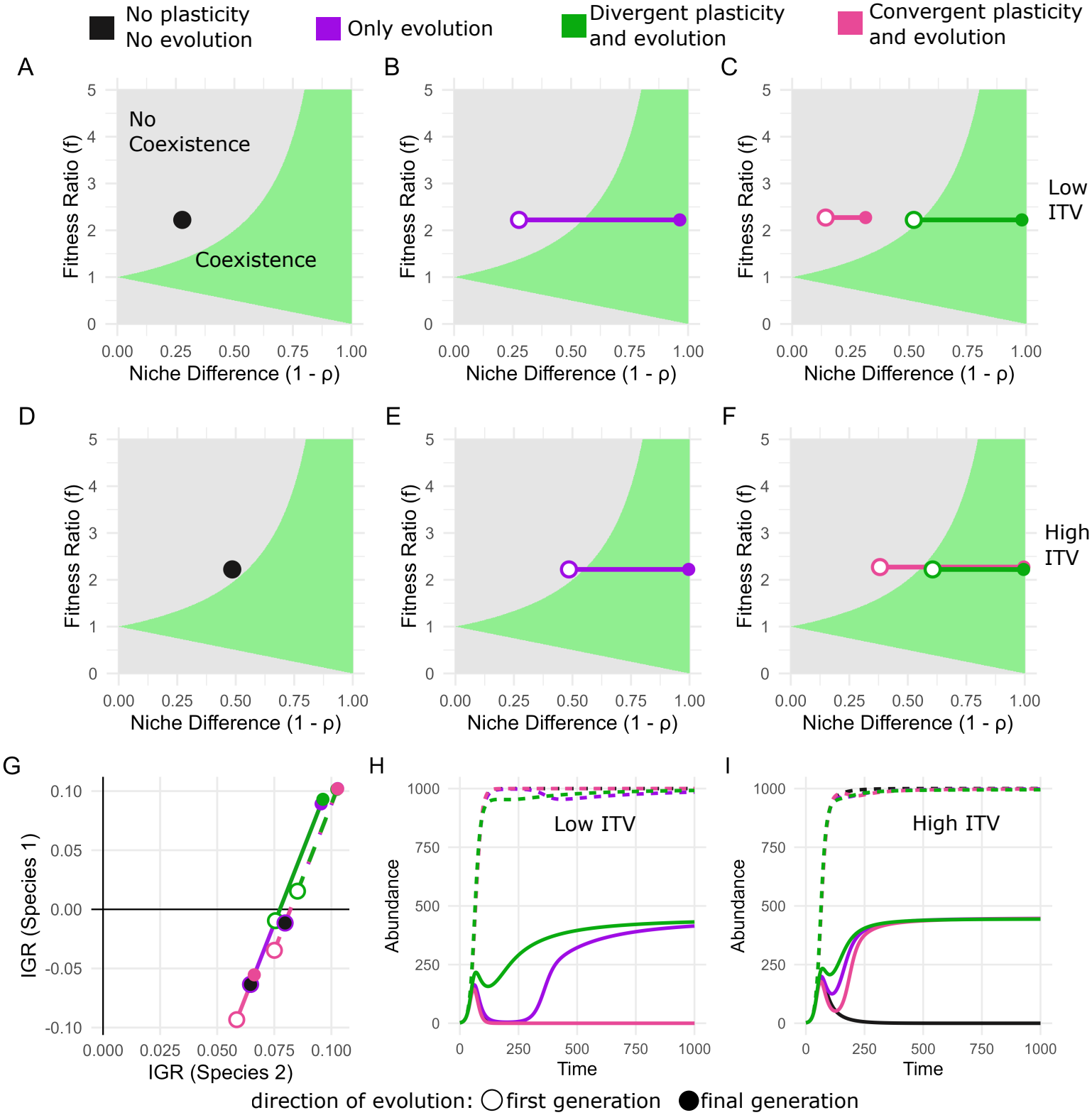
Effect of phenotypic plasticity, evolution and ITV in a broader environment: Light green and grey regions indicate where coexistence is possible or not, respectively. Open circles mark the first generation and closed circles the last generation to indicate the direction of evolutionary change. (**A**-**C**) correspond to low ITV (*σ*_1_ = *σ*_2_ = 0.2), while (**D**-**F**) correspond to high ITV (*σ*_1_ = *σ*_2_ = 0.8). (**G**) shows the evolution of the invasion growth rates of both species with solid lines indicating low ITV and dashed lines indicating high ITV scenarios. The scenarios overlap and fall on the same line and are jittered slightly (*≈*0.003) down-left and up-right, respectively, along both IGR axes for visual clarity. (**H**) and (**I**) show the population dynamics for low ITV and high ITV, respectively, with dashed lines denoting the dominant species (species 2) and solid lines the weaker species (species 1). Parameter values used: *θ* = 6.5, *ω* = 1.1. Other parameter values used are given in Table 1. The detailed dynamics of each of these scenarios are shown in Supplementary figures S2-S9, S18 and S21.

The difference in the starting points in figure 2A-F is due to the effect of plasticity (figure 2 B,C;E,F) within a generation and ITV (figure 2 A,D;B,E;C,F) at the beginning of every generation. For low ITV (first row of figure 2), the system maintained its initial state of competitive exclusion in the absence of plasticity and evolution (figure 2A), but evolved to coexistence when evolution was present (figure 2B). Divergent plasticity accelerated this transition. The increased niche difference (or reduction in the competition coefficient, *α*_12_) (figure S4B compared to figure S3B) caused by divergent plasticity and evolution pushed the invasion growth rate of the weaker species from negative values to positive values in fewer generations (figure S4D compared to figure S3D) and thus evolution to coexistence (green lines in figure 2C) was faster. Contrary to this, convergent plasticity hindered evolution to coexistence, resulting in competitive exclusion (pink lines in figure 2C). In this case, convergent plasticity increased the competition coefficient within a generation (figure S5B compared to figure S3B). Evolution selected for traits that reduced niche overlap in future generations (figure S5A), but this alone was insufficient to prevent extinction. The population-dynamic consequences are shown in figure 2H. Without evolution or plasticity, the weaker species went extinct (black solid line, behind the pink solid line). Evolution prevented extinction, producing the classical U-shaped recovery in population density characteristic of evolutionary rescue (Carlson et al., 2014; Gomulkiewicz & Holt, 1995). Divergent plasticity facilitated evolutionary rescue of the weaker species, whereas convergent plasticity prevented rescue (figure 2H).

Higher ITV, through increased genetic variance, increased the evolutionary pressure due to competition (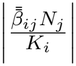 for species *i* in Eq. 9) (figure S1D-F compared to figure S1A-C), resulting in greater trait divergence and lower niche overlap than under low ITV across three plasticity scenarios (figure S7A-B vs S3A-B; figure S8A-B vs S4A-B; figure S9A-B vs S5A-B). This resulted in the high ITV scenarios (figure 2D-F) to be more conducive to coexistence compared to the respective low ITV scenarios (figure 2A-C). Consequently, with only evolution, the system reached the coexistence region faster, divergent plasticity produced coexistence from the first generation, and convergent plasticity delayed but did not prevent coexistence. These differences were also reflected in higher invasion growth rates (figure 2G) and faster evolutionary rescue of the weaker species (figure 2I).

### 3.2 Role of plasticity in a narrower environment

Reduced environmental breadth (*θ*=0.7, *ω*=0.55) imposed an additional constraint on evolution by limiting the amount of trait divergence that is possible without incurring fitness costs. The initial generation of the scenarios shown in the narrow environment (figure 3) is more coexistence-friendly compared to the broader environment scenario, as the competition width here used here is smaller (*ω*=0.55) than in the broader environment case (*ω*=1.1).

**Figure 3:**
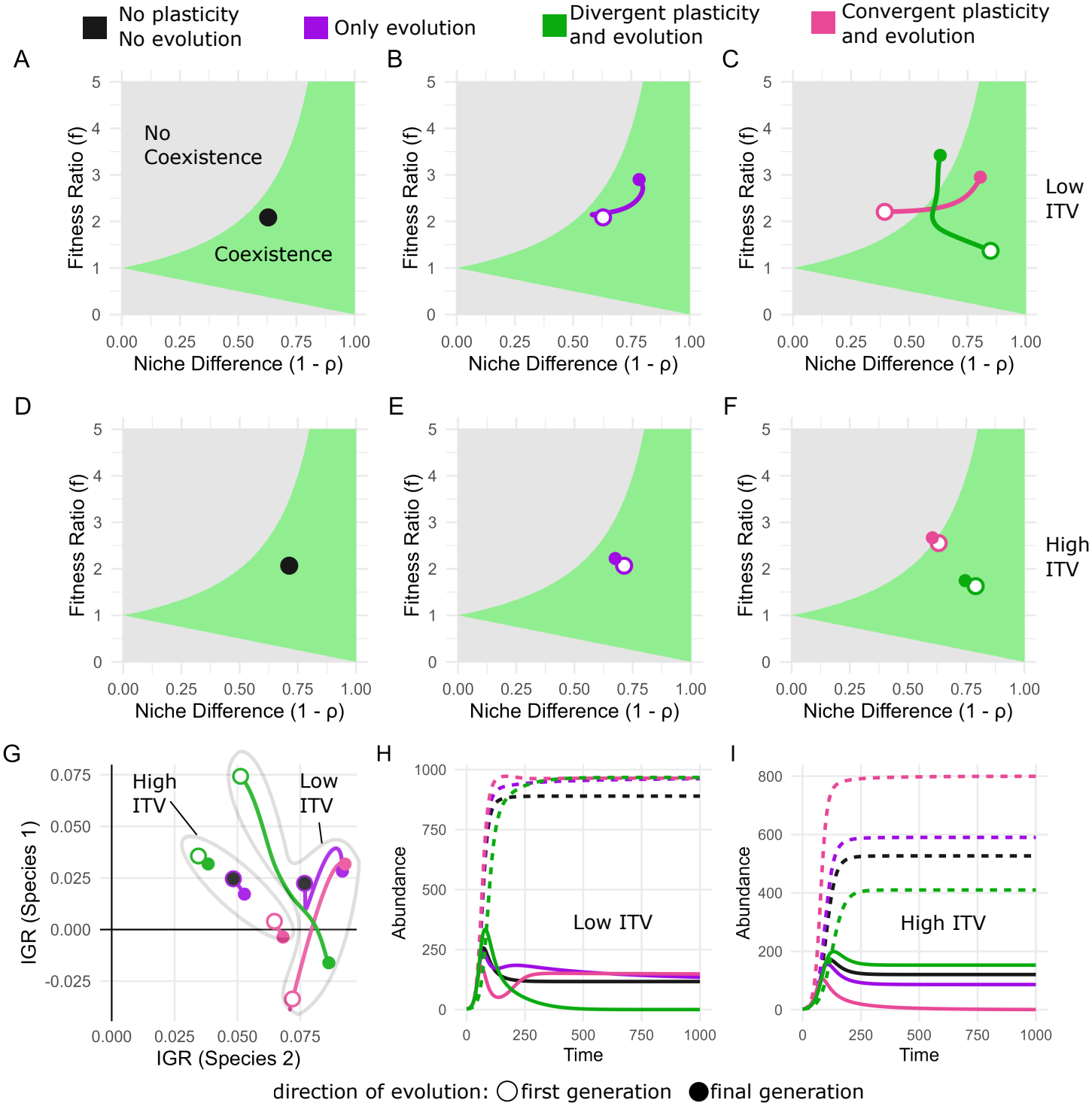
Effect of phenotypic plasticity and evolution in a narrow environment: Light green and grey regions indicate where coexistence is possible or not, respectively. Open circles mark the first generation and closed circles the last generation to indicate the direction of evolutionary change.(**A**-**C**) correspond to low ITV (*σ*_1_ = *σ*_2_ = 0.2), while (**D**-**F**) correspond to high ITV (*σ*_1_ = *σ*_2_ = 0.8). (**G**) shows the evolution of the invasion growth rates of both species with solid lines indicating low ITV and dashed lines indicating high ITV scenarios. The scenarios overlap and are slightly jittered for visual clarity. (**H**) and (**I**) show the population dynamics for low ITV and high ITV, respectively, with dashed lines denoting the dominant species (species 2) and solid lines the weaker species (species 1). Parameter values used: *θ* = 0.7, *ω* = 0.55. Other parameter values used are given in table 1. The detailed dynamics of each of these scenarios are shown in Supplementary (figure S10-S13, figure S19, figure S20, figure S22).

For low ITV, without evolution and plasticity, the system maintained its initial state of coexistence (figure 3A). Evolution was shaped not only by evolutionary pressure to reduce competition but also to maximize growth. We found that 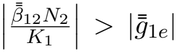 for the weaker species 1 and 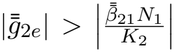 for the dominant species 2 (figure S1G-I). Consequently, the dominant species converged towards the environmentally optimal trait range (i.e., towards [*−θ, θ*]) maximizing its growth, whereas the weaker species diverged away from it reducing competition. The magnitude of convergence (of the dominant species) and divergence (of the weaker species) depended on the respective plasticity scenario (figure S11A, S12A, S13A). With evolution alone, the system evolved deeper into the coexistence region (figure 3B). When divergent plasticity was present, the system initially remained within the coexistence region but ultimately evolved toward competitive exclusion (figure 3C). Divergent plasticity pushed expressed phenotypes beyond the environmentally optimal trait range within a generation (figure S19B), generating selection on dominant species 2 that shifted its mean trait toward more overlapping regions of trait space (figure S19B, figure S12A) compared with evolution alone (figure S11A). This increased population-level competition (figure S12B), increased the intrinsic growth rate of the dominant species, and reduced that of the weaker species (figure S12G). Consequently, the invasion growth rate of the weaker species declined to negative values (figure 3G, figure S12D), leading to its exclusion (figure 3H). In contrast, with convergent plasticity, the system initially laid outside the coexistence region but eventually evolved toward coexistence (figure 3C). Convergent plasticity maintained expressed phenotypes within the environmentally optimal trait range (figure S19C), allowing evolutionary divergence of the mean trait of weaker species 1 (figure S19C, figure S13A). Over evolutionary time, this reduced population-level competition (figure S13B) while limiting reductions in the intrinsic growth rate of the weaker species (figure S13G). Together, these effects increased its invasion growth rate to positive values (figure 3G, figure S13D), resulting in coexistence (figure 3H). Although the weaker species initially showed the highest population growth rate when evolution acted with divergent plasticity and the lowest when evolution acted with convergent plasticity, over evolutionary timescales, divergent plasticity ultimately drove the weaker species to competitive exclusion (figure 3H).

An increased ITV causes a larger fraction of individuals of both species to fall outside the environmental breadth (figure S20) inducing strong evolutionary pressure to increase intrinsic growth rate (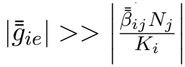 for species *i*, figure (S1J-L)). This caused the species’ trait distributions to converge to the trait value of 0, which is midway between *−θ* and *θ* (figure S15A) to minimize the loss in intrinsic growth rates (figure S15G). Compared to the scenario without evolution (figure 3D), evolution increased niche overlap (figure S15B), but the system still satisfied 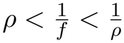 (*≈* positive invasion growth rates both species, figure S15D) to result in coexistence (figure 3E). Divergent plasticity accelerated this convergence (figure S16A) (by pushing the traits outside the environmental breadth, forcing evolution to converge faster) and thereby promoted coexistence (figure 3F,I), whereas convergent plasticity delayed trait convergence (figure S17A), resulting in competitive exclusion (figure 3F,I).

### 3.3 Role of competition width and environment breadth in modulating plasticity-driven eco-evolutionary outcomes

We explored the *θ*–*ω* parameter space to determine the evolutionary outcomes without plasticity (figure 4A). We then examined how divergent (figure 4B) and convergent plasticity (figure 4C) modify these outcomes across the same parameter space. With evolution alone, comparable competition width and environmental breadth (*ω ≈ θ*) marked the transition between coexistence and competitive exclusion (figure 4A) up to a critical competition width (*ω ≈* 1.45), beyond which evolution was not capable of rescuing the weaker species to establish coexistence. Along this boundary (*θ ≈ ω*), divergent plasticity shifted the system toward exclusion (figure 4B), converting parameter combinations that previously supported coexistence into competitive exclusion. In contrast, convergent plasticity shifted regions previously characterized by exclusion to coexistence (figure 4C). At low competition width, we identified a subset of the *θ*-*ω* region in which divergent plasticity (with evolution) restored coexistence from competitive exclusion, whereas convergent plasticity produced the opposite outcome. In a broader environment, divergent plasticity expanded the coexistence regime (figure 4B), while convergent plasticity reduced it (figure 4C).

**Figure 4:**
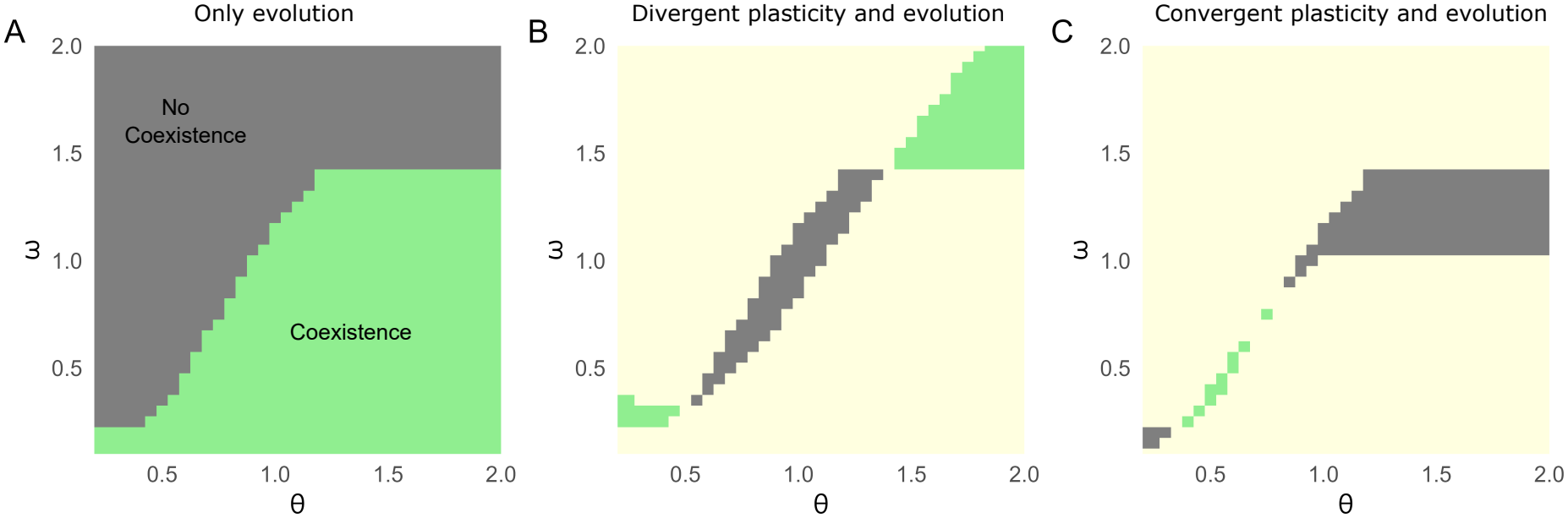
Effect of environmental breadth and competition width (*θ, ω*) in modulating plasticity-driven competition outcomes: Light green and dark grey regions indicate where species coexist or not, respectively. (**A**) Shows the parameter space for which species coexist (and not) with only evolution and no plasticity. *ω≈* 1.45 is the competition width beyond which species cannot coexist. (**B**) Shows how divergent plasticity changes the outcome of (**A**). (**C**) Shows how convergent plasticity changes the outcome of (**A**). In (**B**) and (**C**) light green regions indicate parameter combinations that shift system from competitive exclusion to coexistence, whereas dark grey regions indicate combinations that shift from coexistence to competitive exclusion under divergent plasticity in (**B**) and convergent plasticity in (**C**). In the light yellow region, there is no change of outcome with respect to (**A**). Parameter values used: *K*_1_ = 450, *K*_2_ = 1000, *σ*_1_ = *σ*_2_ = 0.2.

## 4 Discussion

The role of phenotypic plasticity in the eco-evolutionary dynamics of competing species is not well studied. Recent studies tend to decouple plastic (Gibbs et al., 2025; Hess et al., 2022) and evolutionary responses (Hart et al., 2019; Yamamichi et al., 2022) on coexistence. The combined effects of plasticity and evolution on species coexistence have yet to be integrated into a single framework. Here, we provide a theoretical framework which incorporates individual-level plasticity into a generalised Lotka-Volterra eco-evolutionary dynamical model. Our results show that the effect of plasticity on the evolution of coexistence depends on how plasticity changes evolutionary pressures due to competition and growth, and how the resulting evolutionary changes reshape the niche and fitness differences that determine ecological coexistence. Plasticity therefore influences coexistence not just through its immediate effect on niche overlap and fitness differences, but also by altering the subsequent evolution of niche and fitness differences.

In broad environments, intrinsic growth costs were negligible, and evolution was dominated by selection to reduce competition (niche overlap) (figure S1A-F). Divergent plasticity reduced the within-generation niche overlap and facilitated evolution to coexistence (figure 2C,F). Convergent plasticity increased within-generation niche overlap and delayed evolutionary rescue of the weaker species (figure 2C,F,H,I). Under low ITV, this delay was potent enough to result in competitive exclusion (figure 2H), consistent with the idea that plasticity can either buy or reduce time available for evolutionary rescue (Fox et al., 2019; Snell-Rood et al., 2018).

In narrow environments, trait divergence incurred substantial growth costs. Thus, evolution was jointly shaped by evolutionary pressure due to competition and intrinsic growth (figure S1G-L). For lower ITV, divergent plasticity pushed expressed phenotypes beyond the environmentally optimal trait range (i.e., beyond [*−θ, θ*]; figure S19B), redirecting evolution toward greater niche overlap and fitness differences (figure 3C), ultimately excluding the weaker species (figure 3H). In contrast, convergent plasticity maintained expressed phenotypes within the favourable range while allowing inherited mean traits to evolve a reduced niche overlap which balanced the fitness differences (figure S19C), restoring positive invasion growth for the weaker species (figure 3G). Thus, a plastic response that reduces competition and results in coexistence in the short term need not promote coexistence over longer evolutionary timescales.

ITV further shaped coexistence by modifying ecological parameters and the strength of growth- and competition-mediated selection. In broad environments, high ITV reduced niche overlap and strengthened selection for trait divergence, promoting coexistence. In narrow environments, high ITV instead promoted convergence toward the environmental optimum; divergent plasticity accelerated this convergence and facilitated coexistence, whereas convergent plasticity delayed it and caused exclusion of the weaker species. This is especially relevant given recent evidence that competition can drive character convergence rather than divergence (Leonard et al., 2026). Thus, plasticity can determine whether such evolution facilitates or hinders coexistence.

Environmental breadth (*θ*) and competition width (*ω*) determine the evolutionary pathways through which coexistence emerges. Depending on the *θ*-*ω* scenario, under evolution alone, co-existence can arise through divergent evolution of both species (figure S3A), convergence of one species to the center of the environmental range and divergence of the other (figure S11A), or convergence of both species (figure S15A). Plasticity modifies these evolutionary pathways differently along the boundaries between coexistence and exclusion, reversing long-term coexistence outcomes (figure 4B-C). When coexistence is promoted by divergence (broad environment, top right region in figure 4A) or convergence of both species (narrow environment, bottom left region in figure 4A), divergent plasticity expands the coexistence region, whereas convergent plasticity contracts it. In contrast, when coexistence arises because one species evolves towards the environmental optimum while the other evolves away from it (narrow environment, intermediary region in figure 4A), divergent plasticity can contract the coexistence region, whereas convergent plasticity can expand it.

### 4.1 Broader implications, extensions and limitations

Our results show that understanding the evolution of coexistence requires jointly considering both evolution and plasticity. The short-term positive effect of plasticity on coexistence (as in Hess et al., 2022) can be disrupted over evolutionary timescales. Conversely, whether evolution facilitates or disrupts coexistence (as in Pastore et al., 2021) can be altered by plasticity.

Contributing to the debate on whether plasticity facilitates or hinders evolution, we show that its effects can be asymmetric: the same plastic response may slow evolution in one species while accelerating it in its competitor (see Supplementary section S1.7). Whether plasticity is adaptive or maladaptive is less straightforward in the context of species interactions where each species reciprocally forms the selective environment for the other (see Supplementary section S1.8).

We model plasticity as continuous trait dynamics, extending beyond standard reaction norms along environmental gradients (Dupont et al., 2024) and treatments as binary switches (Bennett et al., 2023) or stage-structured frameworks (Brass et al., 2021). Although previous theoretical frameworks have examined how abiotic plasticity interacts with evolution to affect species persistence (Chevin & Chauhan, 2025; Chevin et al., 2010), none have incorporated biotic plasticity in shaping these outcomes. Our model accounts for the type of plasticity by incorporating density-dependent divergent and convergent plasticity, consistent with recent empirical studies of competition-induced plasticity (Bebout et al., 2026).

Several assumptions in our framework point to directions for future work. We held plasticity coefficients constant, preventing the evolution of plasticity. We did not incorporate abiotic plasticity (Chevin & Chauhan, 2025; Chevin et al., 2010), which, along with biotic plasticity, could jointly shape eco-evolutionary dynamics. Our framework could be extended to multi-species communities to help understand how plasticity influences community structure (Miner et al., 2005). We did not consider conspecific-induced plasticity, nor spatially localised responses (Brown et al., 2026; Gibbs et al., 2026). Finally, we assumed constant phenotypic variance. Relaxing this and allowing trait distributions to evolve may reveal additional feedbacks between ecological and evolutionary processes.

## 5 Acknowledgements

This work was funded by German research foundation (DFG) as part of research consortium SFB-TRR 212, project numbers 316099922 and 396782288.

## 6 Author Contributions

V.V. and M.W. conceptualized the study. G.B. contributed to the conceptual development of the study. V.V. designed the methodology with input from M.W., developed the computational model, conducted the analyses, prepared the visualizations and wrote the original manuscript. All authors reviewed and edited the manuscript drafts.

## S1 Supplementary Information

### S1.1 Derivation for the within-generation plasticity-driven trait dynamics

With *t* denoting within-generation time and *T* the number of the current generation, let *N*_1_(*T* ) and *N*_2_(*T* ) be the population densities of species 1 and 2 in a generation. 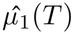 and 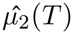 are the baseline inherited mean trait of each species at the beginning of the generation and which are not affected by the competition-induced phenotypic plasticity with a generation. *µ*_1_(*t*) and *µ*_2_(*t*) are the mean trait of the species affected by phenotypic plasticity at a time t within generation. Thus, 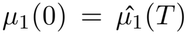 and 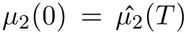. The dynamics of the trait of the *i*^*th*^ individual of species 1 are given by

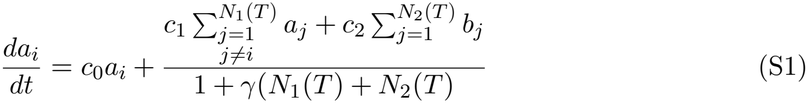

and the dynamics of the *i*^*th*^ individual of species 2 are

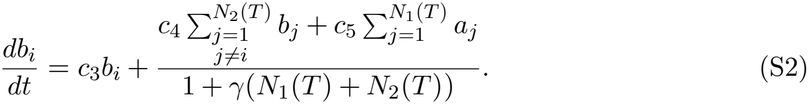

Here, the trait of each individual of each species is linearly dependent on its own trait, traits of all conspecific individuals and all heterospecific individuals. We know

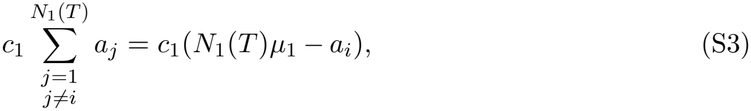

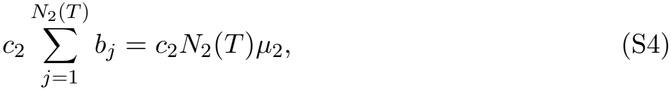

and

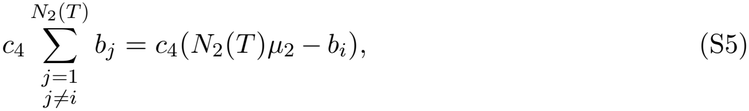

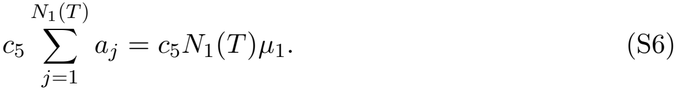

Substituting Eq. S3, Eq. S4, Eq. S5 and Eq. S6 in Eq. S1 and Eq. S2, respectively, we get

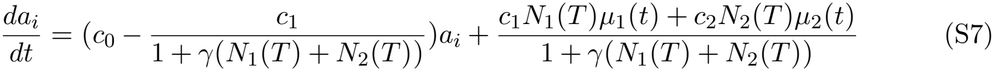

and

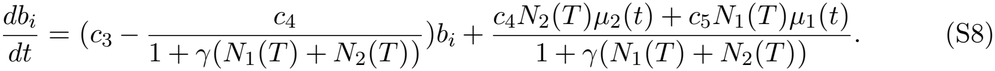

With each individual trait *a*_*i*_ and *b*_*i*_ dynamically changing due to biotic phenotypic plasticity, the corresponding mean traits of the phenotypic trait distribution (*µ*_1_(*t*) and *µ*_2_(*t*)) also change. We can derive the analytical expression for the mean trait dynamics as follows:

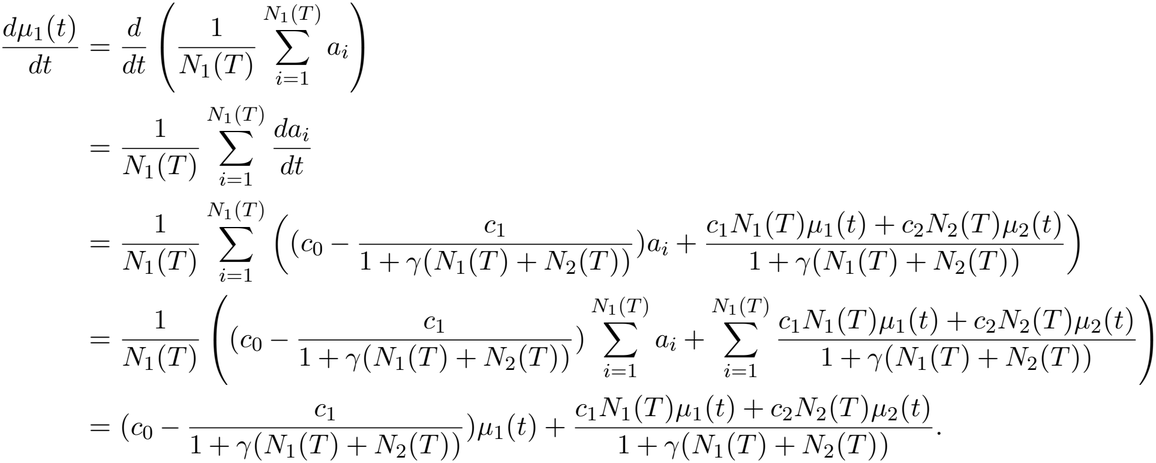

Similarly, the mean trait dynamics of species 2 are given by

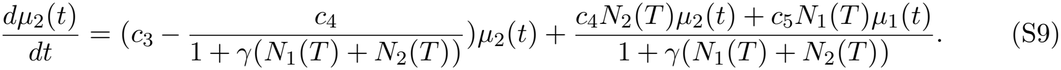

The mean trait dynamics are in the form of linear coupled differential equations. We can analytically solve this using the matrix diagonalization method. For ease of notation, we subsume the trait saturation factor into the plasticity coefficients:

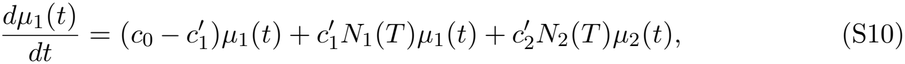

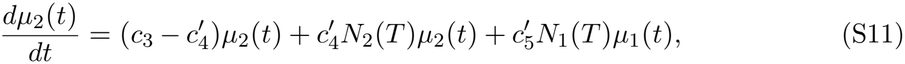

where 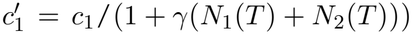 and analogously for the other coefficients. Since we consider only heterospecific plasticities, we consider *c*_1_ = *c*_4_ = 0. The mean trait dynamics now read:

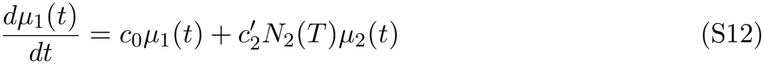

and

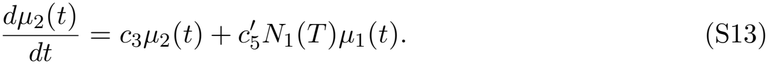

In matrix form, we can rewrite these expressions as

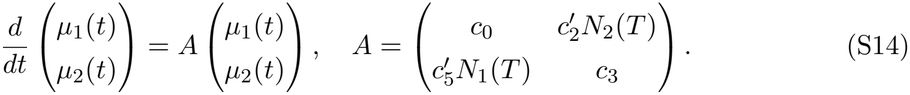

For coupled differential equations of this form, we can obtain analytical solutions which have the form

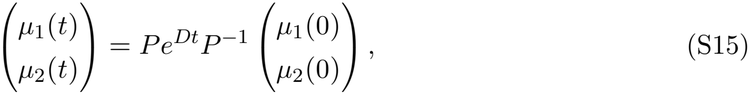

where *P* is the matrix of eigenvectors of *A, D* is the corresponding diagonal matrix of eigenvalues and *P* ^*−*1^ is the inverse of *P*. For this, we find the eigenvalues *λ*_1_ and *λ*_2_ of the coefficient matrix *A* as given by

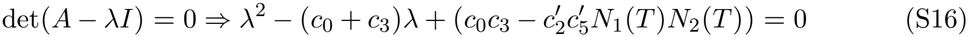

This gives,

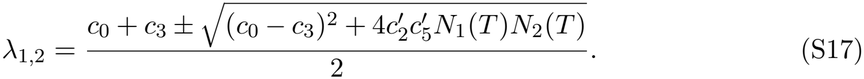

To obtain eigenvectors *v*_*i*_ for each eigenvalue, for each *λ*_*i*_, solve (*A − λ*_*i*_*I*)***v***_*i*_ = 0. This gives eigenvectors as

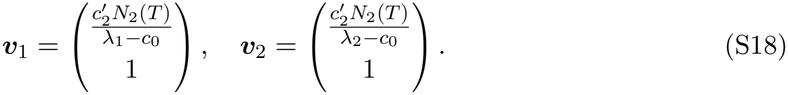

Thus

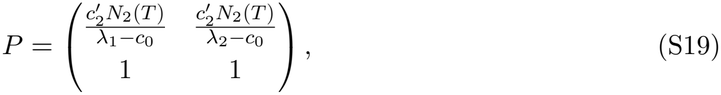

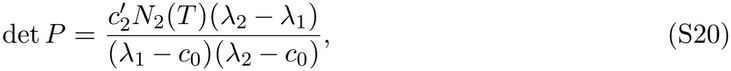

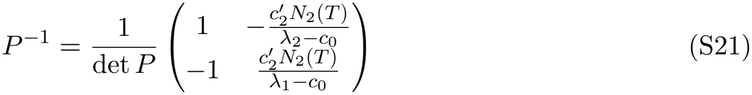

and

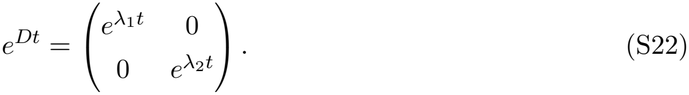

This gives,

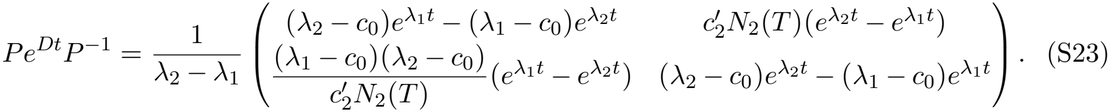

From the characteristic equation Eq. S16, we have

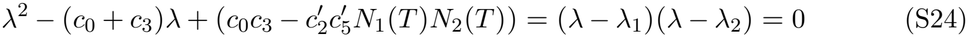

because *λ*_1_ and *λ*_2_ are the roots of this quadratic equation. Evaluating this expression at *λ* = *c*_0_ gives

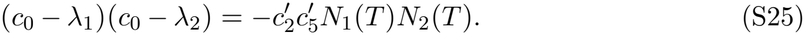

Thus, S23 becomes,

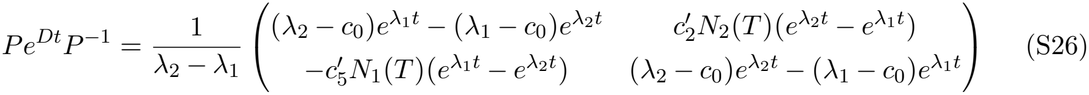

and with

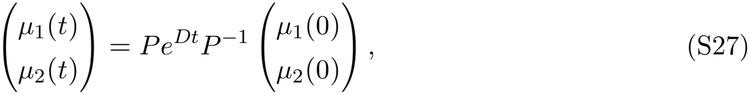

we get

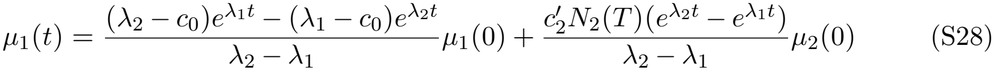

and

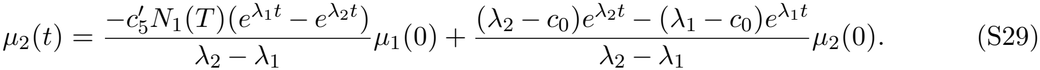

From Eqs. S28 and S29, the within-generation mean trait dynamics can be written as,

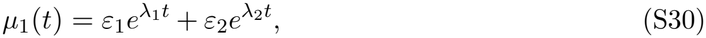

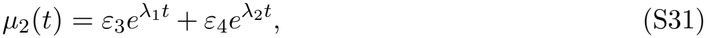

where

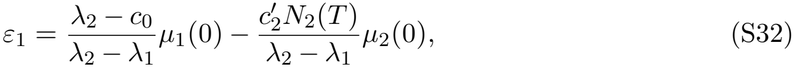

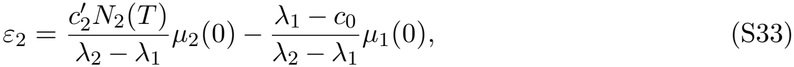

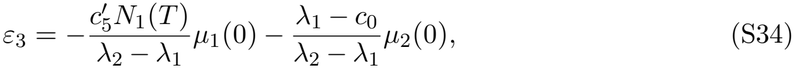

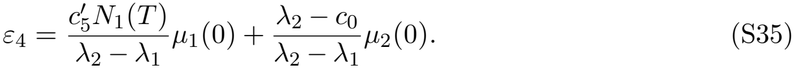

Substituting Eq. S31 in Eq. S7 with the assumption that *c*_1_ = 0, we get

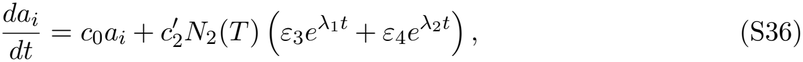

or equivalently,

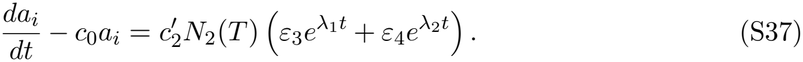

This is a first-order linear differential equation of the form 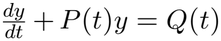 that can be solved using the integrating factor method. Its integrating factor is

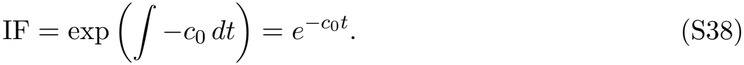

Multiplying both sides by the integrating factor gives

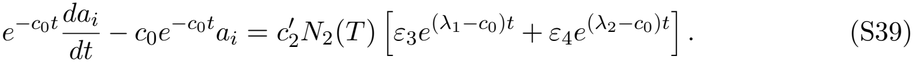

Since

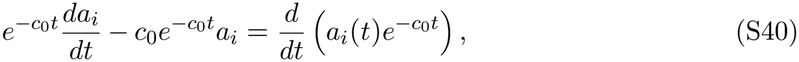

we obtain

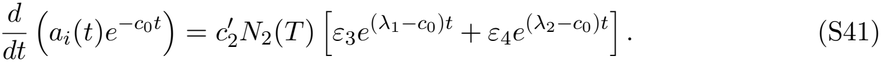

Integrating Eq. S41 from 0 to *t*, we get

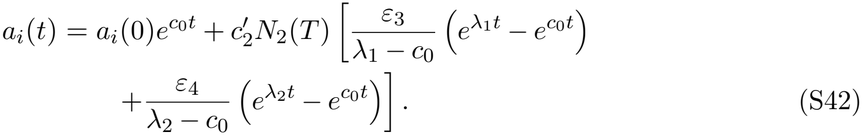

Similarly, substituting Eq. S30 in Eq. S8 with the assumption that *c*_4_ = 0, we get

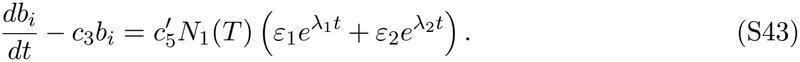

The integrating factor in this case is

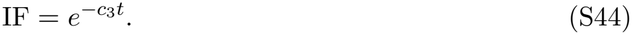

Multiplying Eq. S43 throughout by the integrating factor and rearranging, we get

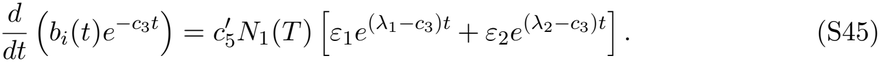

Integrating Eq. S45 from 0 to *t*, we get

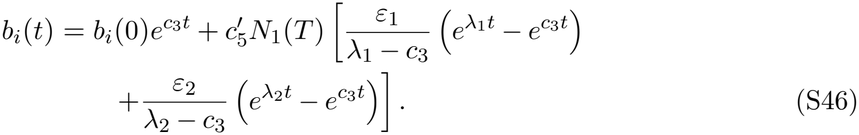

Substituting 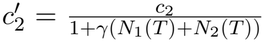 and 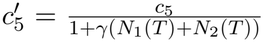 in Eq. S42 and Eq. S46 respectively we get expression for plasticity subsumed trait value of species 1 and 2 as

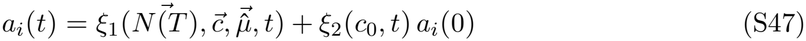

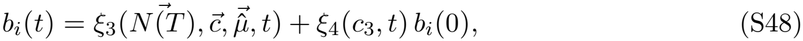

where

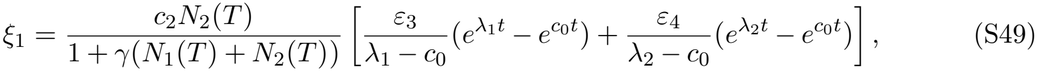

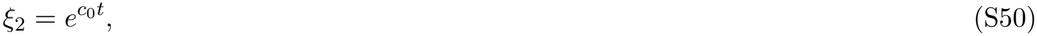

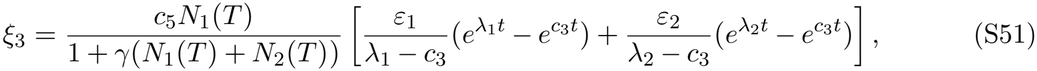

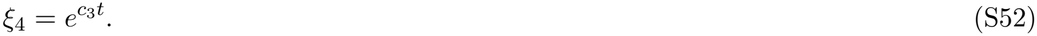

and

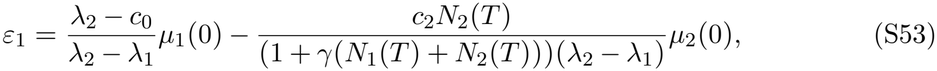

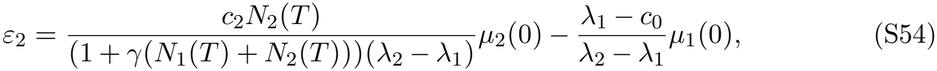

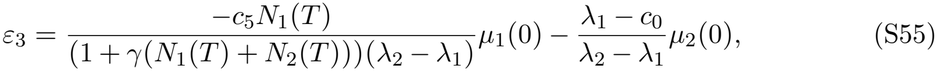

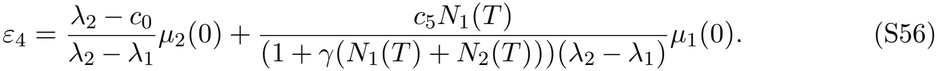

### S1.2 Eco-evolutionary dynamics

#### S1.2.1 Ecological Dynamics

With the within-generation trait dynamics of individuals from each species given by Eq. S47 and Eq. S48, we now use the following two-species Lotka-Volterra population dynamics equations to approximate the growth of population densities of both species from one generation to next.

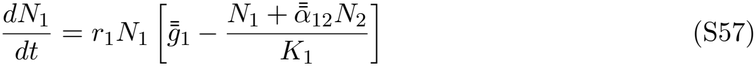

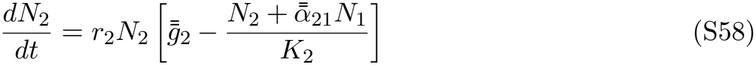

Here, 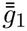 and 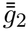 are the species-level intrinsic growth rates of species 1 and 2, respectively. 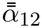 and 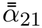 are the species-level interspecific competition coefficients (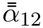is the competition effect on species 1 due to species 2 and 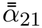 is that on species 2 due to species 1). We assume the intraspecific competition coefficients to be 1. *K*_1_ and *K*_2_ are the variables that dictate the carrying capacities of species 1 and 2, respectively.

##### Intrinsic growth rate

At the individual level, the intrinsic growth is assumed to be a rectangular growth function Eq. S59 having a uniform value of 1 when the trait of the individual falls within the bound of [-*θ, θ*], where *θ* is the environment breadth. This takes the form

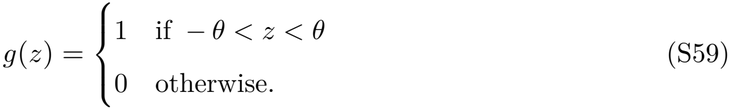

When individual traits, which are *a*(0) and *b*(0) at the beginning of a generation, adjust themselves due to phenotypic plasticity as given by Eq. S47 and Eq. S48, the intrinsic growth function S59 undergoes transformation as given by

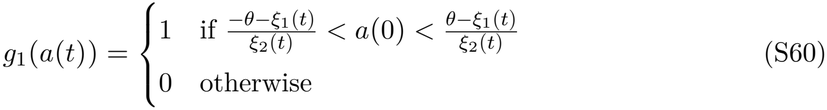

and

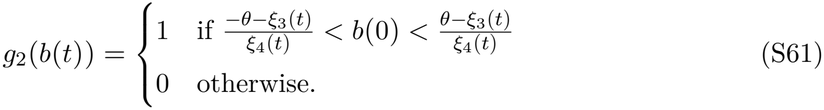

Species-level intrinsic growth rates can then be obtained by nonlinearly averaging these individual-level expressions temporally over the generational time *τ* and the trait space of the respective species as given by

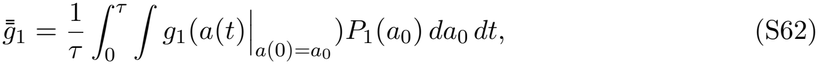

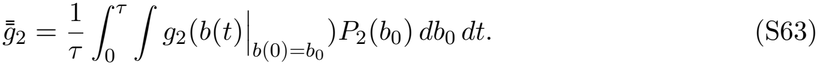

##### Competition coefficients

At the individual level, we consider a Gaussian interaction kernel to capture the effect of competition as given by

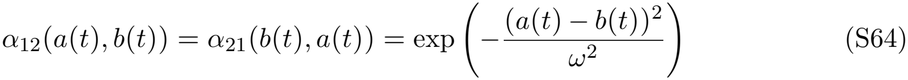

where *ω* is the competition width. At the species level, when each individual adjusts its trait plastically, the competition coefficient can be nonlinearly averaged across the generational time and the trait space as follows,

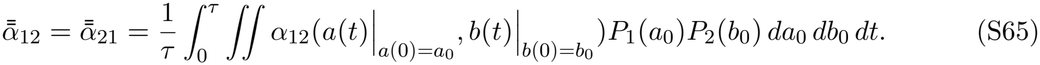

This serves as the within-generation competition coefficient subsuming the effects of individual trait variation and competition-induced plasticity.

#### S1.2.2 Evolutionary Dynamics

We use the framework of Barabás & D’Andrea (2016) to derive the evolutionary equations. The derivations are given in Supplementary section S1.2.3. The dynamics of mean traits of each species are given by

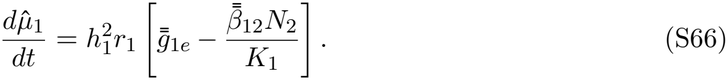

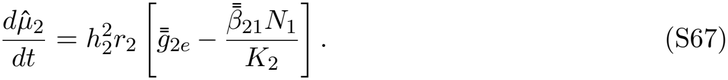

Here 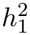 and 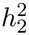 are the heritabilities of species 1 and 2. 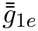 and 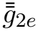 are the species-level evolutionary pressures due to their own growth. 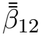 is the evolutionary pressure on species 1 due to competition with species 2 and 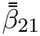 is the evolutionary pressure on species 2 due to competition with species 1. Thus, equations Eq. S57, Eq. S58, Eq. S66 and Eq. S67 together give the eco-evolutionary dynamics of our model.

To incorporate the competition-induced plasticity, we use the framework of Barabás & D’Andrea (2016) (see their supplementary section 2) to find the expressions for evolutionary pressures due to growth and competition. With individuals of species adjusting their traits plastically in response to the heterospecifics, we can get species-level evolutionary pressures due to own growth for species 1 and 2 as given by

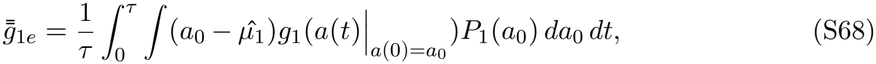

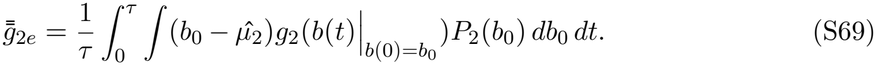

Similarly, with individuals plastically adjusting their traits in response to the heterospecifics, at the species level, the evolutionary pressure on each species due to competition by the other species can be expressed as

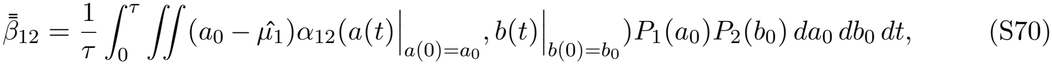

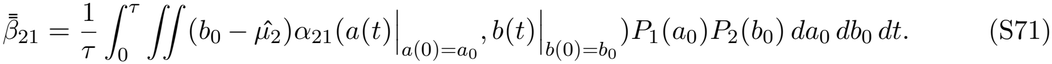

Using the analytical expressions for plastically modified individual traits in Eq. S47 in the competition kernel (Eq. S64), the average species-level competition coefficient at a withingeneration time *t* (i.e., 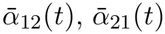) can be obtained by nonlinearly averaging *α*_12_(*a*(*t*), *b*(*t*)) (= *α*_21_(*b*(*t*), *a*(*t*))) over the trait space as

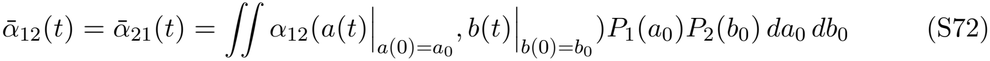

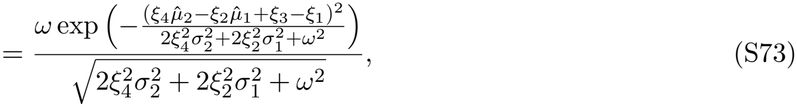

where *ξ*_1_, *ξ*_2_, *ξ*_3_ and *ξ*_4_ are given by Eq. S49, Eq. S50, Eq. S51 and Eq. S52 respectively. Similarly, the expression for the average species-level evolutionary pressures due to competition 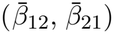 at a within-generation time *t* can be obtained by nonlinearly averaging over the trait space as

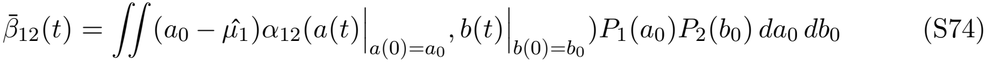

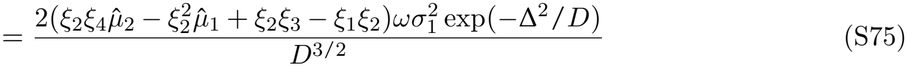

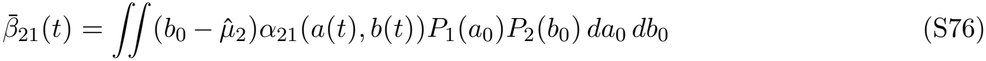

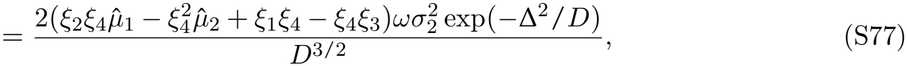

where

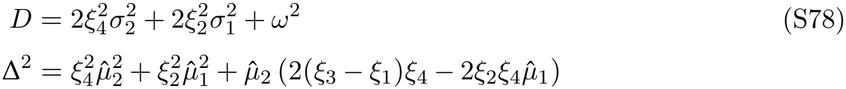

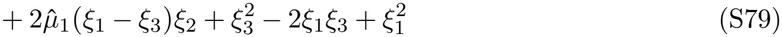

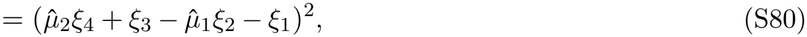

and *ξ*_1_, *ξ*_2_, *ξ*_3_ and *ξ*_4_ are given by Eq. S49, Eq. S50, Eq. S51 and Eq. S52 respectively. Analytical expressions of Eq. S73, Eq. S75, Eq. S77 are obtained using wxMaxima (Maxima, 2023).

#### S1.2.3 Derivation of quantitative genetic Lotka-Volterra model

Let *t ∈* [0, *τ* ] denote within-generation time and *T* denote the between-generation time. The per-capita growth rate of an individual of species 1 with trait *a*_0_ at within-generation time *t* is given by

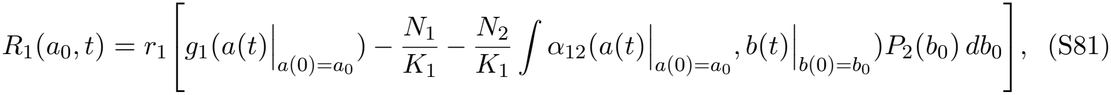

where *a*(*t*) and *b*(*t*) are the plastically modified trait values of individuals initially having traits *a*_0_ and *b*_0_, respectively. *P*_2_(*b*_0_) is the normally distributed trait distribution of species 2 with mean 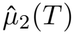 and standard deviation *σ*_2_.

The plasticity-subsumed average per-capita growth rate of an individual of species 1 with initial trait *a*_0_ is given by integrating Eq. S81 over generation time *τ* as

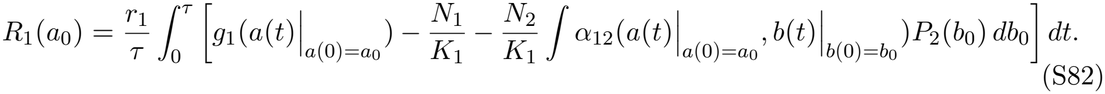

This gives,

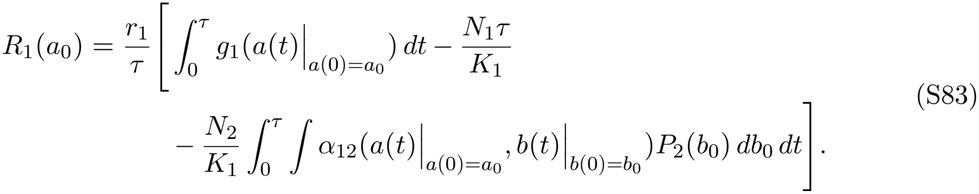

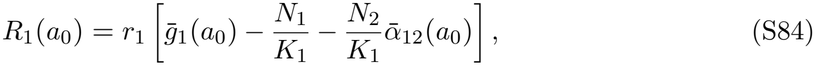

where 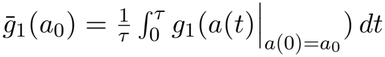 and 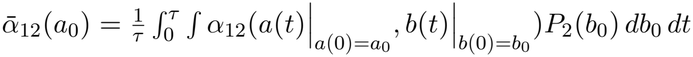.

Population growth is then obtained by averaging the resulting individual growth rate over the trait space as

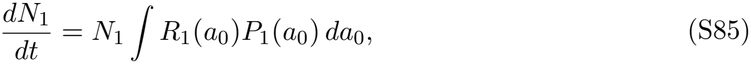

to which substituting Eq. S83 gives

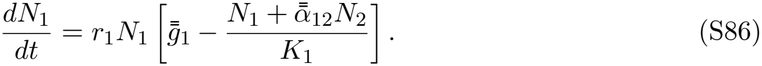

Here we use *P*_1_(*a*_0_) *da*_0_ = 1 and the definitions of 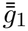 and 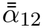 in Eqs. S62 and S65.

Under the quantitative-genetic assumptions, using the framework of Barabás & D’Andrea (2016), we get the evolutionary response as

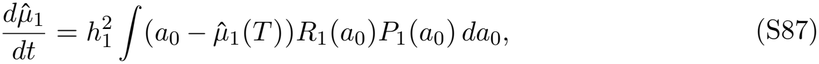

to which substituting S83 gives

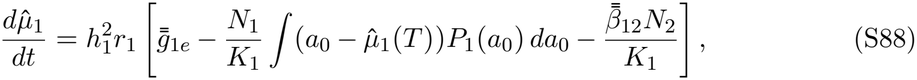

where the evolutionary pressure due to intrinsic growth and interspecific competition terms are defined in Eqs. S68 and S70. Because 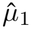 is the mean trait and *P*_1_(*a*_0_) is the normally distributed trait distribution of species 1 with mean 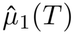 and standard deviation *σ*_1_,

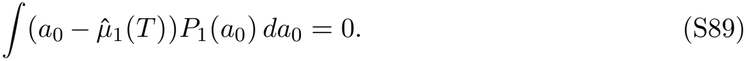

Thus, trait-independent intraspecific competition makes no direct contribution to selection (so there is no 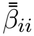 term for species *i*), giving

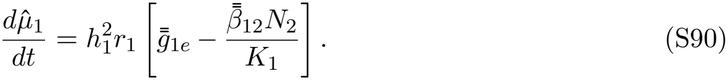

Applying the same steps to species 2 gives Eqs. S58 and S67.

### S1.3 Invasion growth rates

The per capita growth rate of each species is given by,

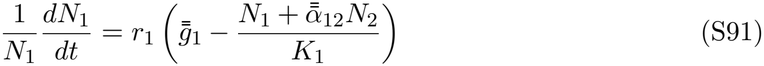

and

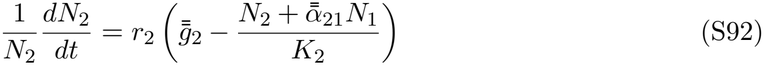

For calculating the invasion growth rate (IGR) of species 2, we assume *N*_1_ at equilibrium 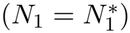 and *N*_2_ *≈* 0 in Eq. S91. From Eq. S91, we get:

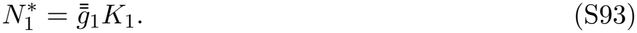

Substituting Eq. S93 in Eq. S92, we obtain the IGR of species 2:

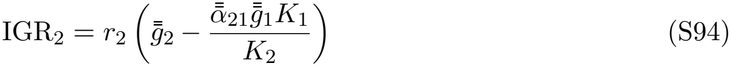

Similarly, for species 1:

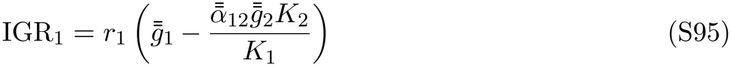

From Eq. S94 and Eq. S95, we see that IGR depends on 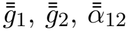, and 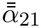. These terms are affected by both evolution and phenotypic plasticity and thus serve as the factor that dictates coexistence of the species.

### S1.4 Niche difference and fitness difference

For the Lotka–Volterra model of the type

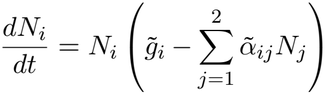

where *N*_*i*_ is species *i*’s population density, *t* is time, 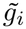 is *i*’s intrinsic growth rate, and 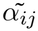 is the competitive effect of species *j* on *i*, the expression for niche overlap (*ρ*) and fitness difference (*f*) for a two-species interaction is as follows (Chesson, 2018; Pastore et al., 2021)

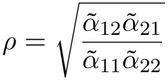

and

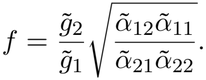

For the Lotka–Volterra equations we have used in our model, Eq. S57 and Eq. S58, we have 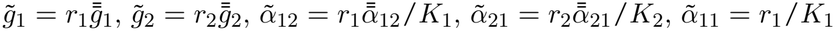*and* 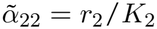.

Thus, the expressions for niche overlap (*ρ*), niche difference (1 *− ρ*) and fitness difference (*f*) are

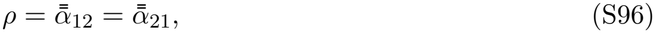

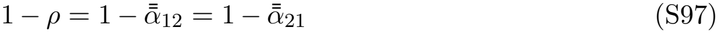

and

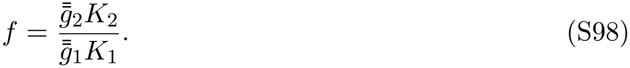

### S1.5 Stability

The Jacobian of the ecological part of the system is given by

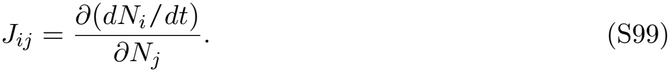

The equilibrium abundances are given by equating Eq. S57 and Eq. S58 to 0, which gives

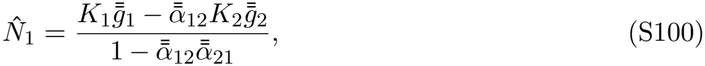

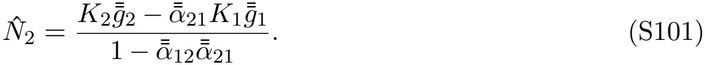

When *J*_*ij*_ is evaluated at equilibrium abundances we get the community matrix of the form

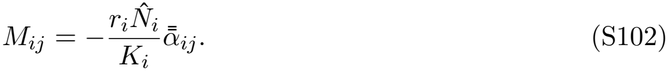

With 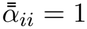, we get,

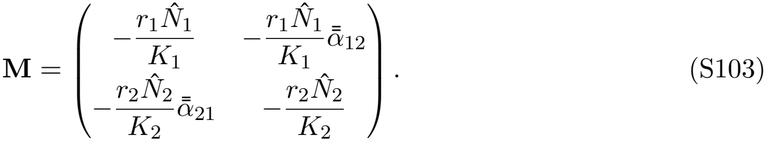

Ecological stability is determined by the real part of the dominant eigenvalue of **M**. The coexistence equilibrium is locally stable when

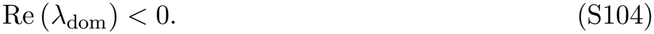

Thus, from Eq. S103 and Eq. S104, competition-induced plasticity affects ecological stability through its effects via the competition coefficients i.e., 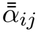 (see Eq. S65) and intrinsic growth rates i.e., 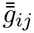 (see Eq. S62, Eq. S63).

### S1.6 Supplementary Figures

**Figure S1:**
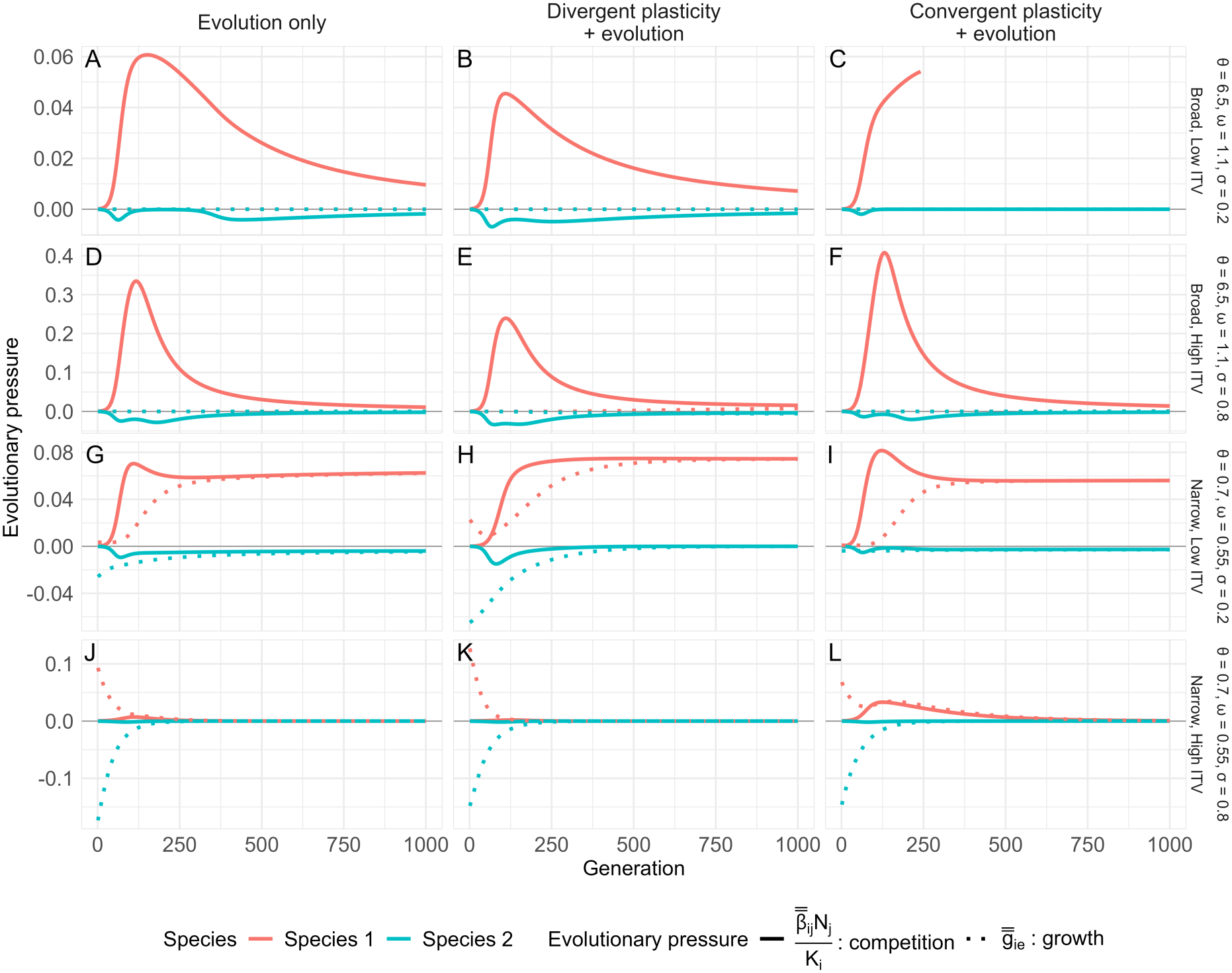
Dynamics of evolutionary pressures: The red solid line shows the evolutionary pressure due to competition for species 1 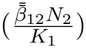. The green solid line shows the evolutionary pressure due to competition for species 2 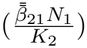. The red dotted line shows the evolutionary pressure due to growth for species 1 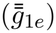. The green dotted line shows the evolutionary pressure due to growth for species 2 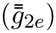. **A-C** shows three plasticity scenarios with broad environment and low ITV, **D-F** shows the scenarios with broad environment and high ITV, **G-I** shows scenarios with narrow environment and low ITV, and **J-L** shows the scenarios with narrow environment and high ITV. We see that evolutionary pressure changes gradually from being competition-driven to intrinsic-growth-driven with environment shifting from broad to narrow.

**Figure S2:**
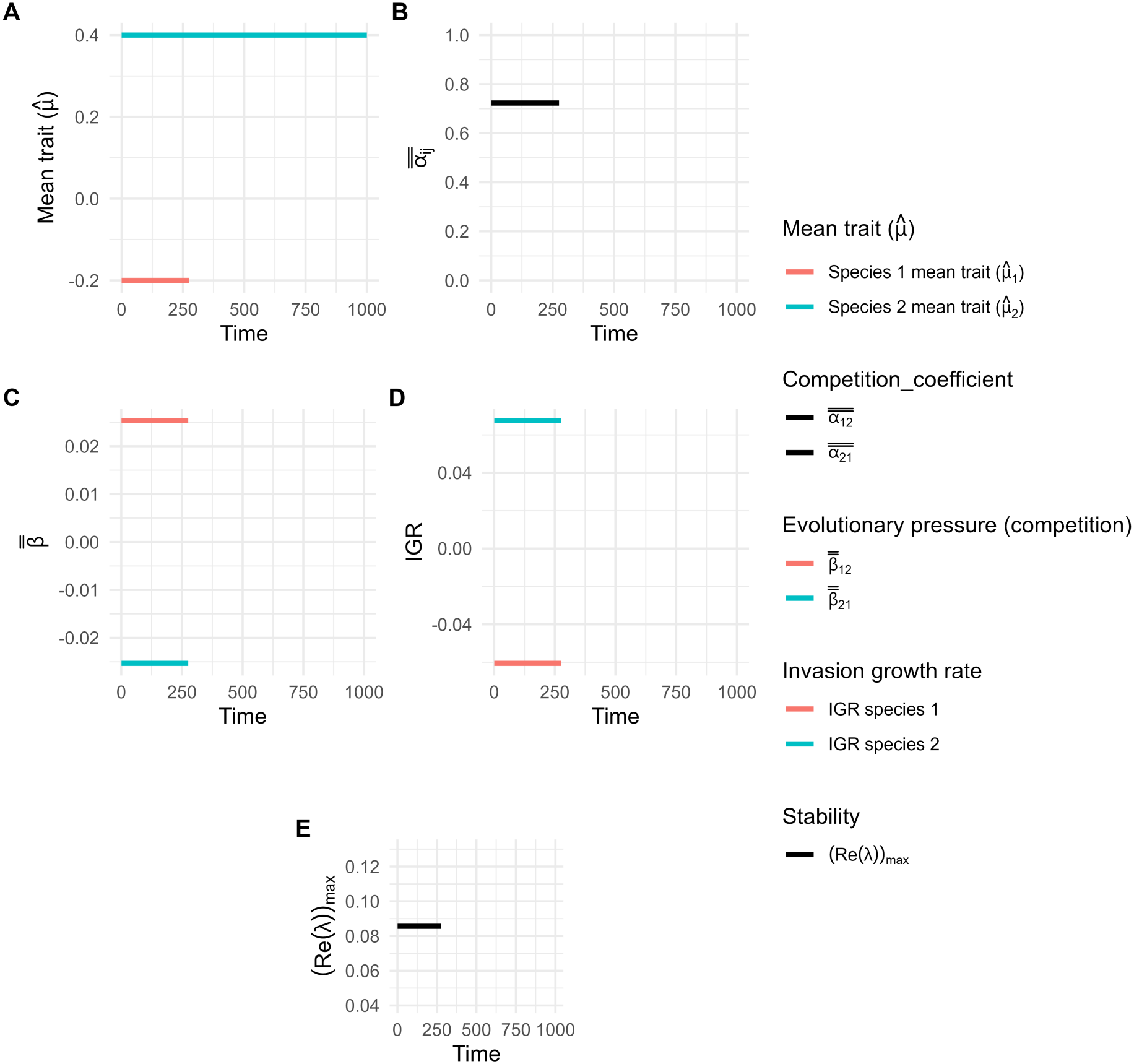
Detailed dynamics for figure 2A-No plasticity, no evolution with low ITV in a broader environment: (**A**) Dynamics of mean traits. (**B**) Dynamics of the competition coefficient (= niche overlap). (**C**) Dynamics of evolutionary pressure due to competition. (**D**) Dynamics of invasion growth rate. (**E**) Dynamics of the dominant eigenvalue of the community matrix (a positive value denotes instability equivalent to no coexistence for a 2-species system; see supplementary section S1.5). The time *t* (abscissa) represents the generations.

**Figure S3:**
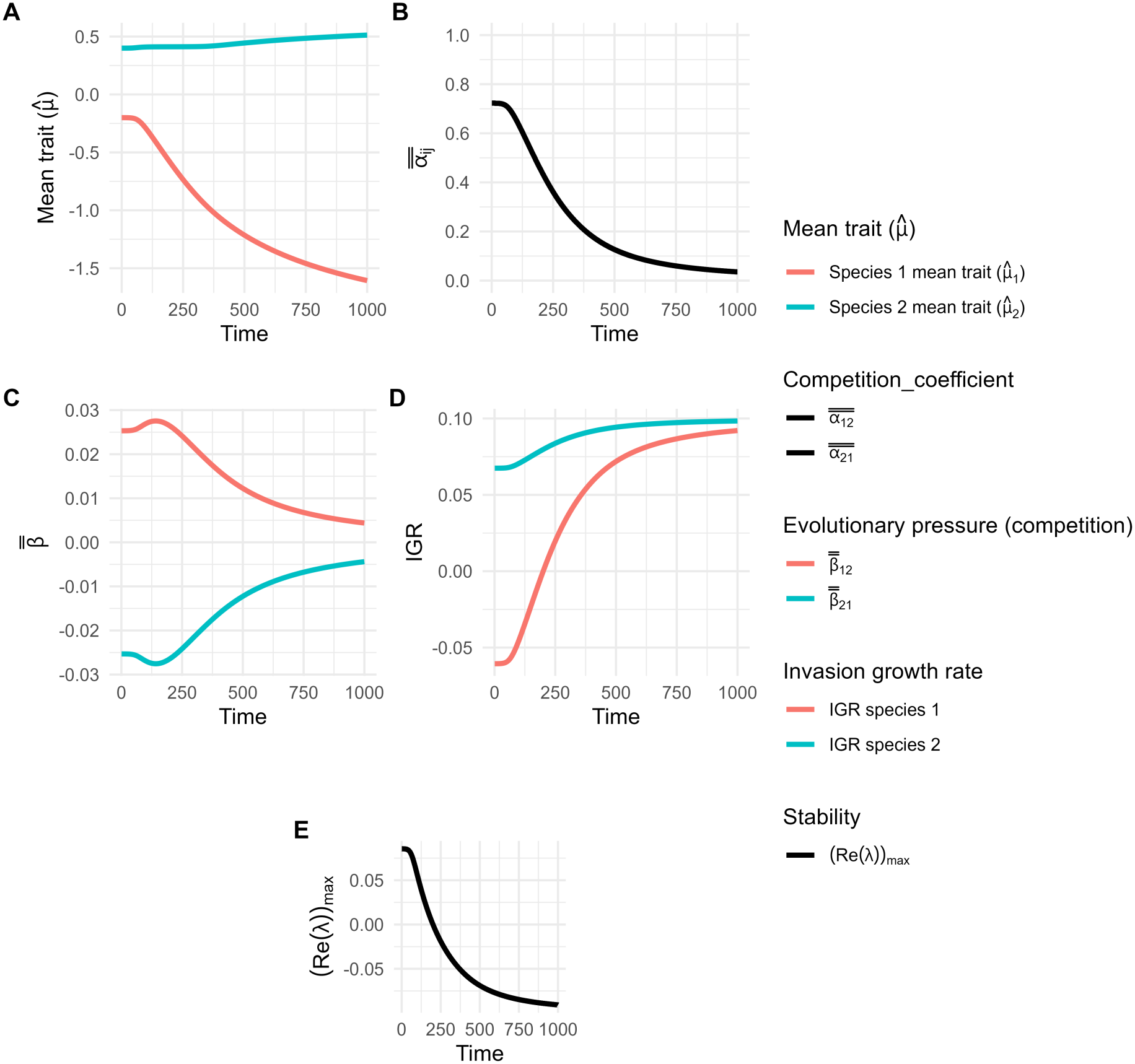
Detailed dynamics for figure 2B-No plasticity, Only evolution with low ITV in a broader environment: (**A**) Dynamics of mean traits. (**B**) Dynamics of competition coefficient (= niche overlap). (**C**) Dynamics of evolutionary pressure due to competition. (**D**) Dynamics of invasion growth rate. (**E**) Dynamics of the dominant eigenvalue of the community matrix (a positive value denotes instability equivalent to no coexistence for a 2-species system; see supplementary section S1.5). The time *t* (abscissa) represents the generations.

**Figure S4:**
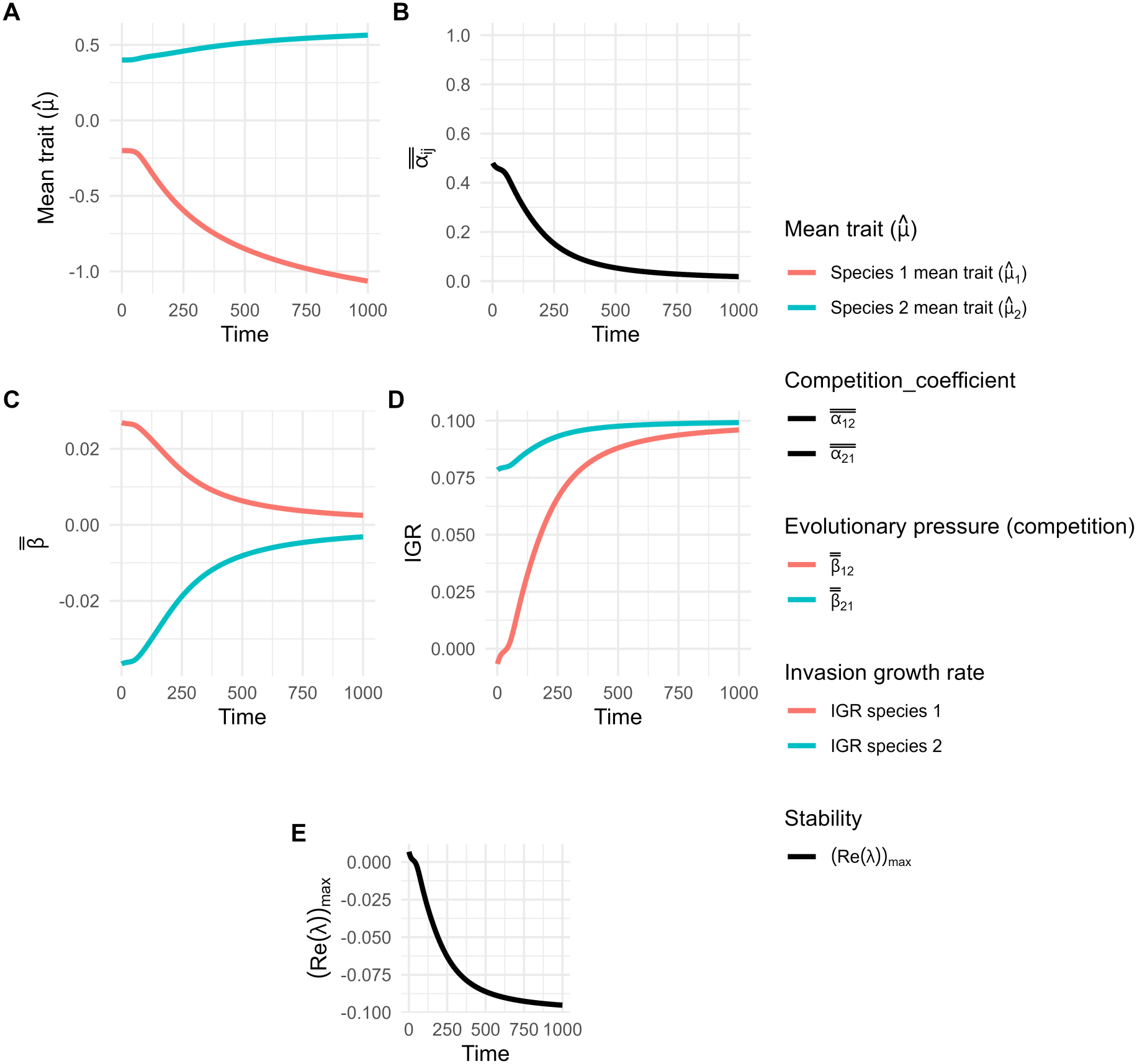
Detailed dynamics for figure 2C-Evolution with divergent plasticity with low ITV in a broader environment: (**A**) Dynamics of mean traits. (**B**) Dynamics of competition coefficient (= niche overlap). (**C**) Dynamics of evolutionary pressure due to competition. (**D**) Dynamics of invasion growth rate. (**E**) Dynamics of the dominant eigenvalue of the community matrix (a negative value denotes stability equivalent to coexistence for a 2-species system; see supplementary section S1.5). The time *t* (abscissa) represents the generations.

**Figure S5:**
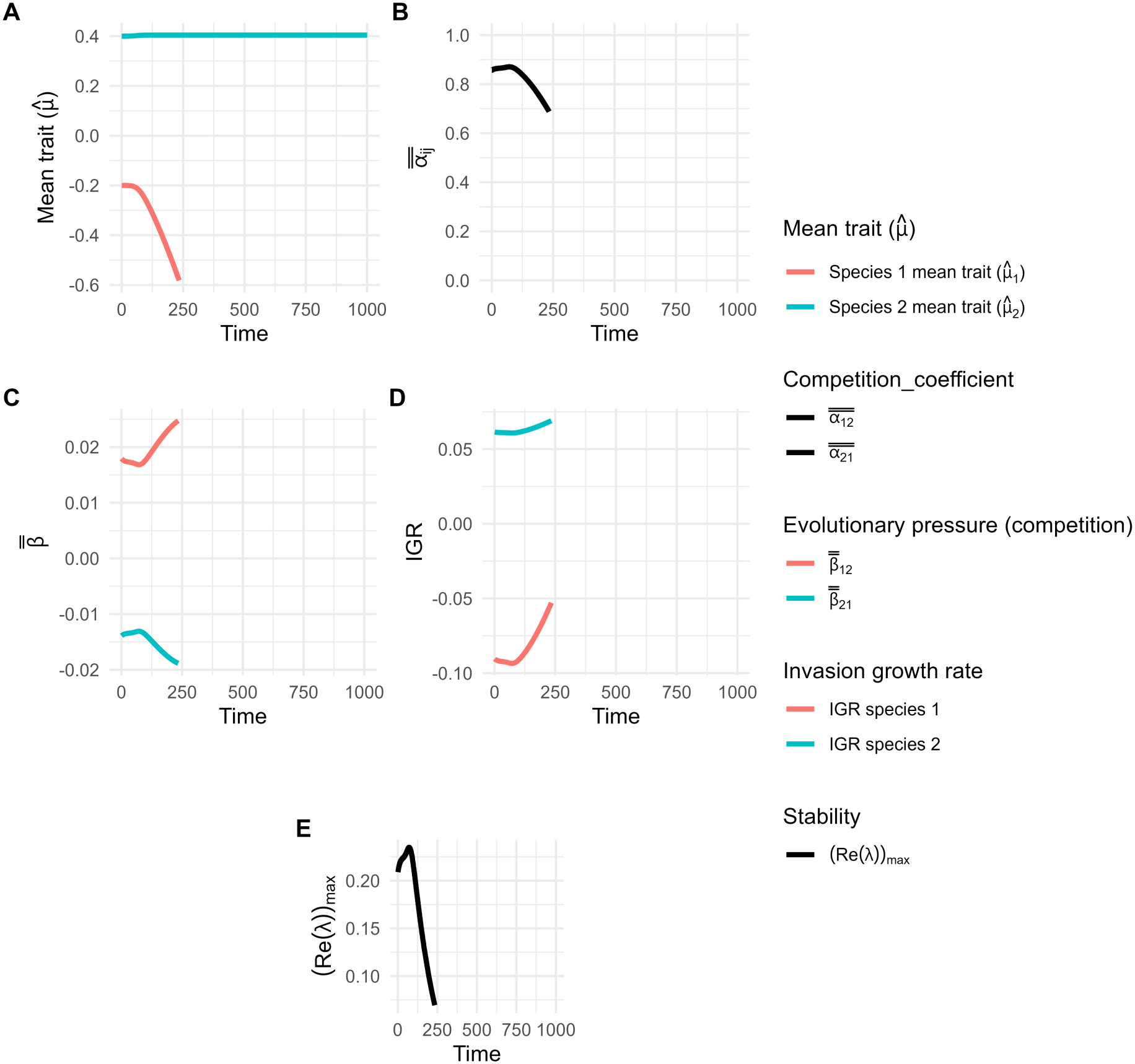
Detailed dynamics for figure 2C-Evolution with convergent plasticity with low ITV in a broader environment: (**A**) Dynamics of mean traits. (**B**) Dynamics of competition coefficient (= niche overlap).(**C**) Dynamics of evolutionary pressure due to competition. (**D**) Dynamics of invasion growth rate. (**E**) Dynamics of the dominant eigenvalue of the community matrix (a positive value denotes instability equivalent to no coexistence for a 2 species system; see supplementary section S1.5). The time *t* (abscissa) represents the generations.

**Figure S6:**
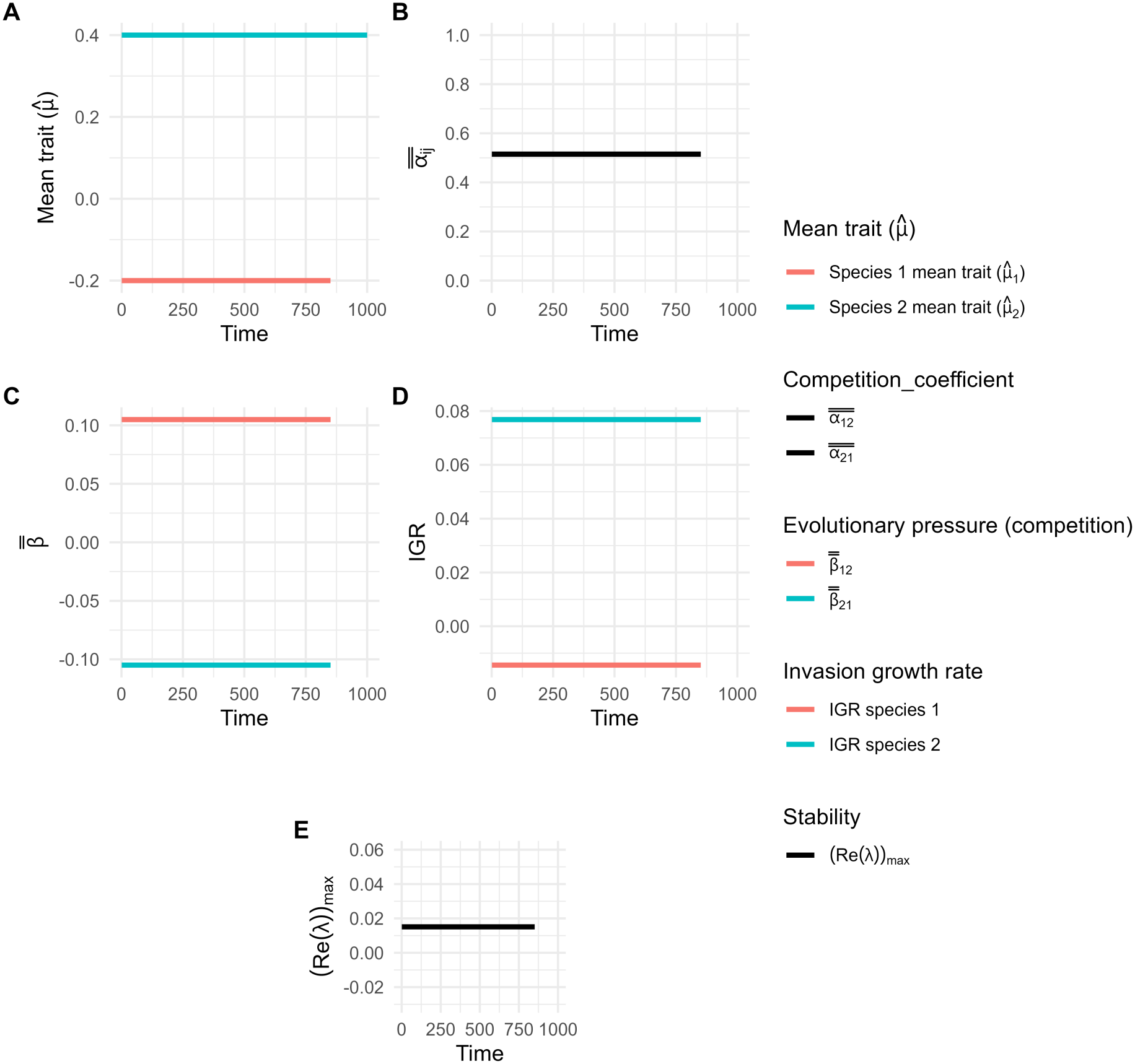
Detailed dynamics for figure 2D-No plasticity, No evolution with high ITV in a broader environment: (**A**) Dynamics of mean traits. (**B**) Dynamics of competition coefficient (= niche overlap). (**C**) Dynamics of evolutionary pressure due to competition. (**D**) Dynamics of invasion growth rate. (**E**) Dynamics of the dominant eigenvalue of the community matrix (a positive value denotes instability equivalent to no coexistence for a 2 species system; see supplementary section S1.5). The time *t* (abscissa) represents the generations.

**Figure S7:**
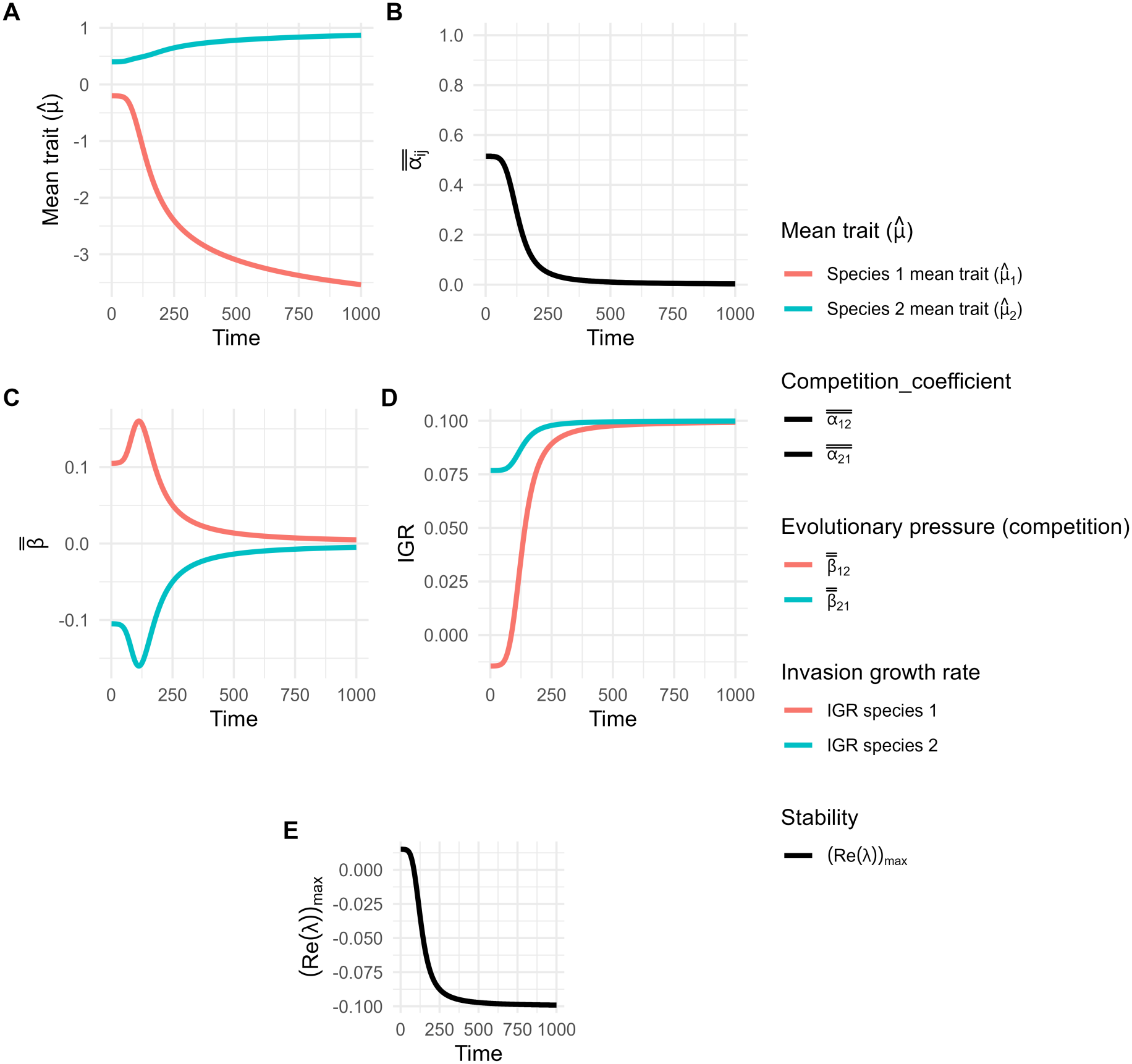
Detailed dynamics for figure 2E-No plasticity, Only evolution with high ITV in a broader environment: (**A**) Dynamics of mean traits. (**B**) Dynamics of competition coefficient (= niche overlap). (**C**) Dynamics of evolutionary pressure due to competition. (**D**) Dynamics of invasion growth rate. (**E**) Dynamics of the dominant eigenvalue of the community matrix (a negative value denotes stability equivalent to coexistence for a 2 species system; see supplementary section S1.5). The time *t* (abscissa) represents the generations.

**Figure S8:**
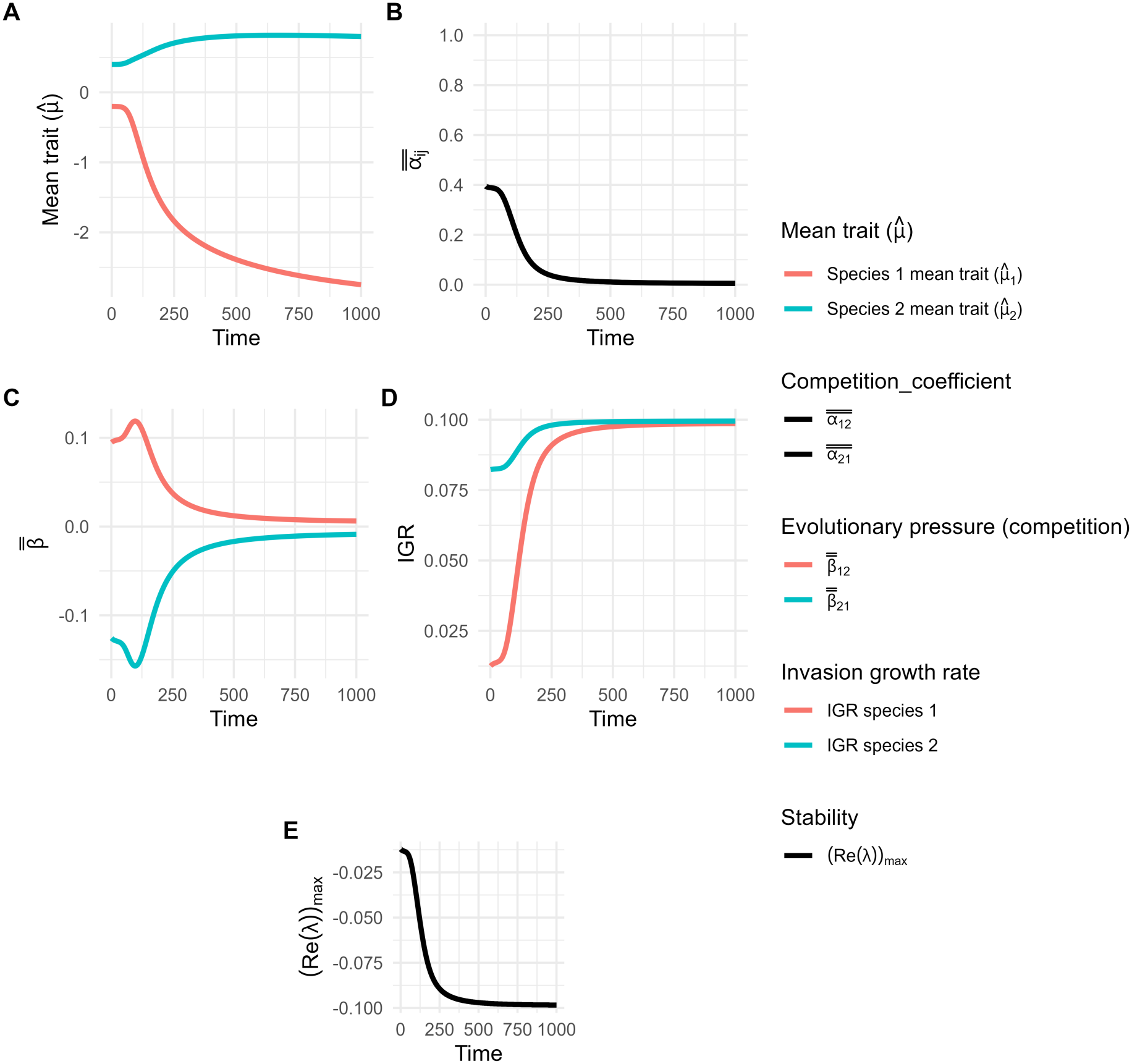
Detailed dynamics for figure 2F-Evolution with divergent plasticity with high ITV in a broader environment: (**A**) Dynamics of mean traits. (**B**) Dynamics of competition coefficient (= niche overlap). (**C**) Dynamics of evolutionary pressure due to competition. (**D**) Dynamics of invasion growth rate. (**E**) Dynamics of the dominant eigenvalue of the community matrix (a negative value denotes stability equivalent to coexistence for a 2 species system; see supplementary section S1.5). The time *t* (abscissa) represents the generations.

**Figure S9:**
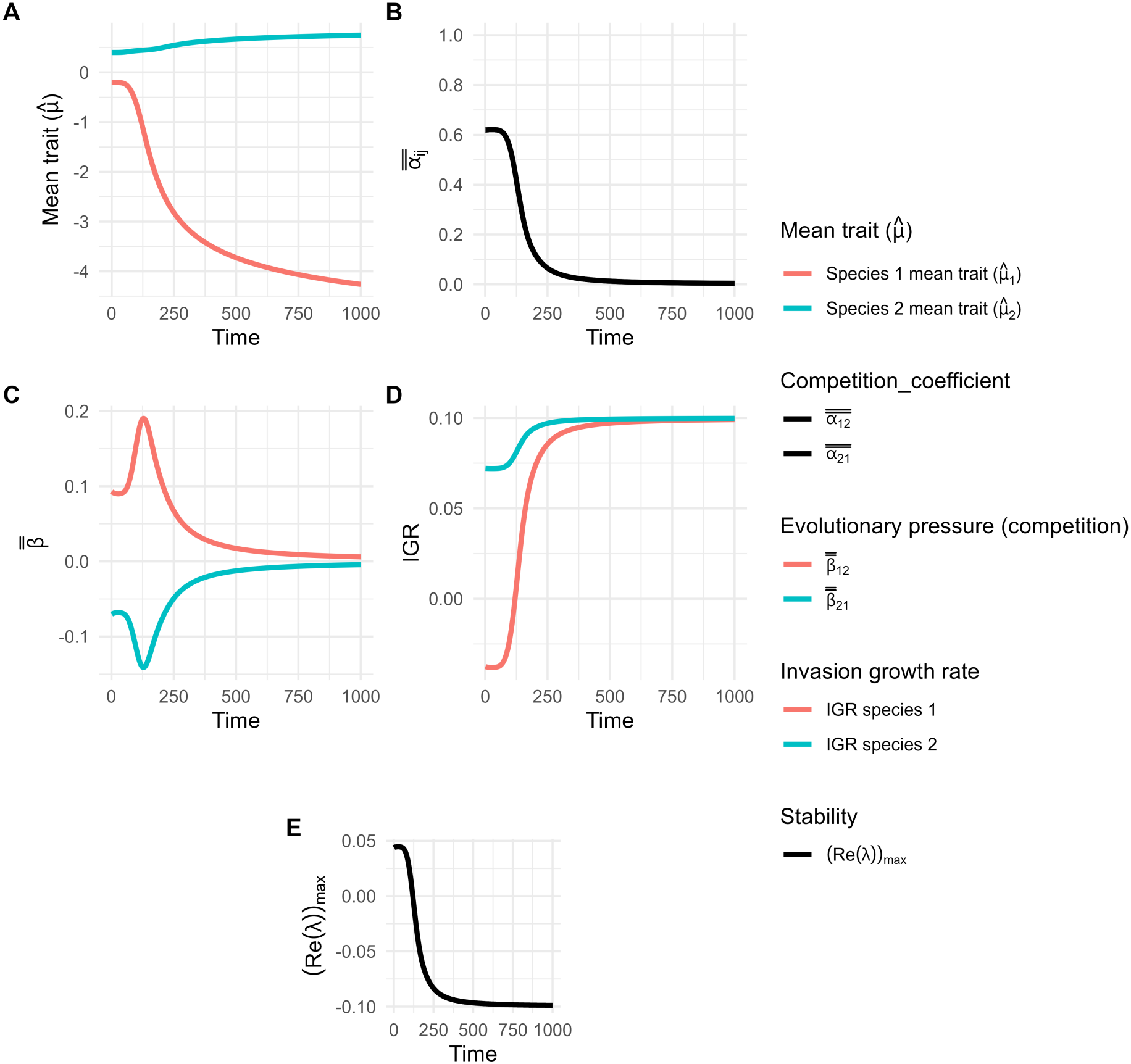
Detailed dynamics for figure 2F-Evolution with convergent plasticity with high ITV in a broader environment: (**A**) Dynamics of mean traits. (**B**) Dynamics of competition coefficient (= niche overlap). (**C**) Dynamics of evolutionary pressure due to competition. (**D**) Dynamics of invasion growth rate.(**E**) Dynamics of the dominant eigenvalue of the community matrix (a negative value denotes stability equivalent to coexistence for a 2 species system; see supplementary section S1.5). The time *t* (abscissa) represents the generations.

**Figure S10:**
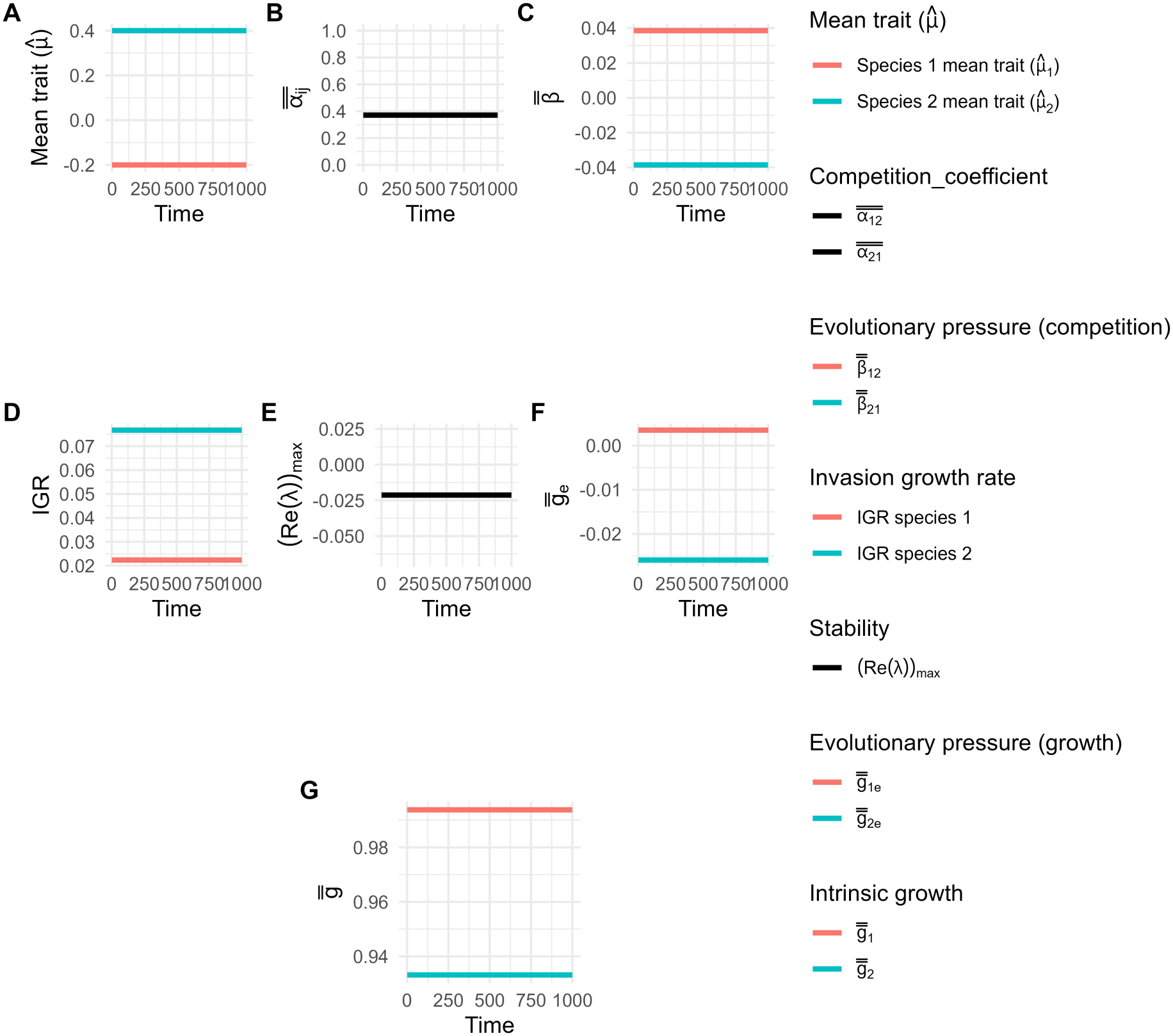
Detailed dynamics for figure 3A-No plasticity and evolution in a narrow environment: (**A**) Dynamics of mean traits. (**B**) Dynamics of competition coefficient (= niche overlap). (**C**) Dynamics of evolutionary pressure due to competition. (**D**) Dynamics of invasion growth rate. (**E**) Dynamics of the dominant eigenvalue of the community matrix (a negative value denotes stability equivalent to coexistence for a 2 species system; see supplementary section S1.5). (**F**) Dynamics of the evolutionary pressure due to growth.(**G**) Intrinsic growth rate. The time *t* (abscissa) represents the generations.

**Figure S11:**
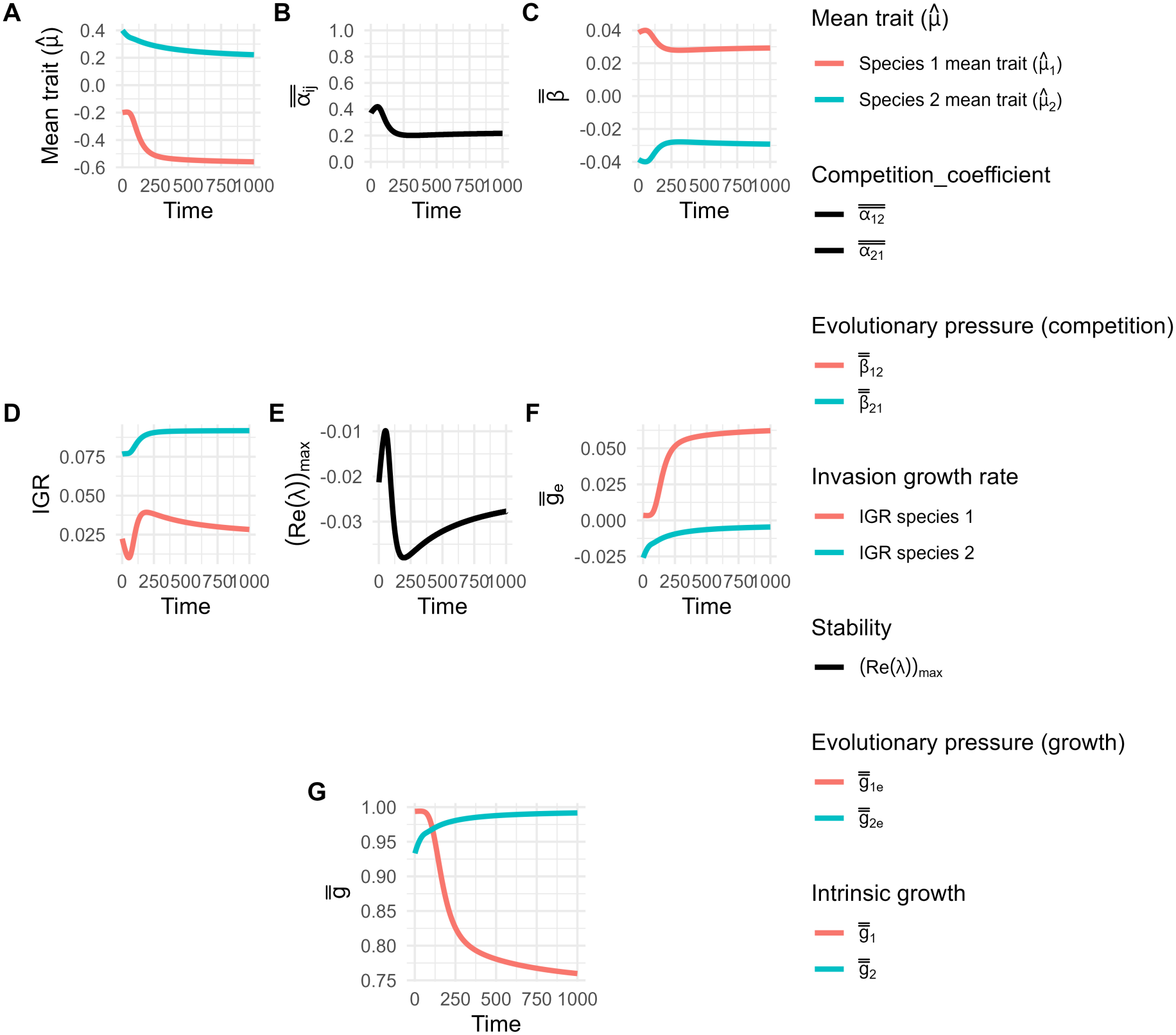
Detailed dynamics for figure 3B-Only evolution in narrow environment: (**A**) Dynamics of mean traits. (**B**) Dynamics of competition coefficient (=niche overlap). (**C**) Dynamics of evolutionary pressure due to competition. (**D**) Dynamics of invasion growth rate. (**E**) Dynamics of the dominant eigenvalue of the community matrix.(Negative value denotes stability equivalent to coexistence for a 2 species system; see supplementary section S1.5). (**F**) Dynamics of the evolutionary pressure due to growth. (**G**) Intrinsic growth rate. The time *t* (abscissa) represents the generations.

**Figure S12:**
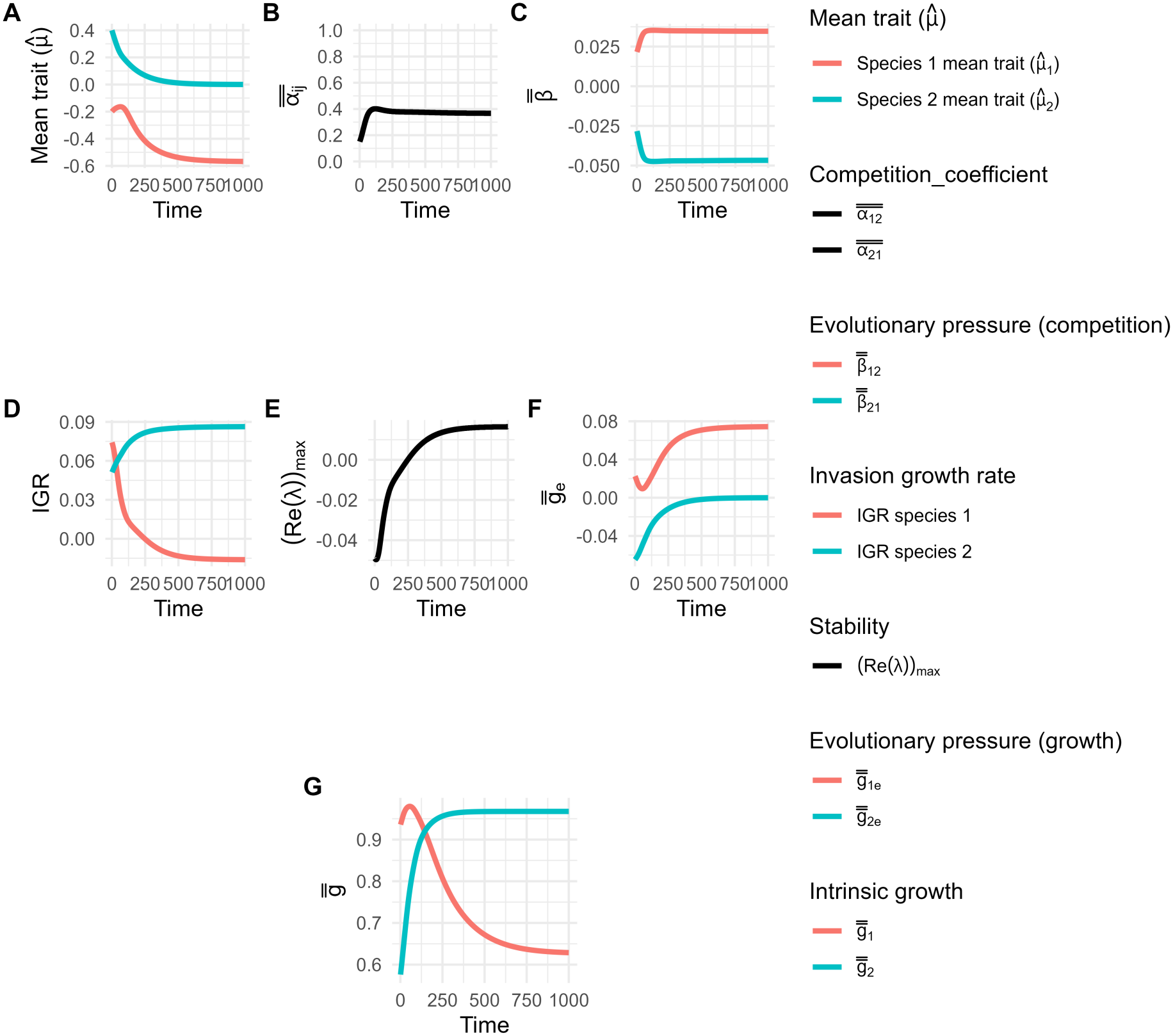
Detailed dynamics for figure 3C-Evolution with divergent plasticity in a narrow environment: (**A**) Dynamics of mean traits. (**B**) Dynamics of competition coefficient (=niche overlap). (**C**) Dynamics of evolutionary pressure due to competition. (**D**) Dynamics of invasion growth rate. (**E**) Dynamics of the dominant eigenvalue of the community matrix.(Positive value denotes instability equivalent to no coexistence for a 2 species system; see supplementary section S1.5).(**F**) Dynamics of the evolutionary pressure due to growth. (**G**) Intrinsic growth rate. The time *t* (abscissa) represents the generations.

**Figure S13:**
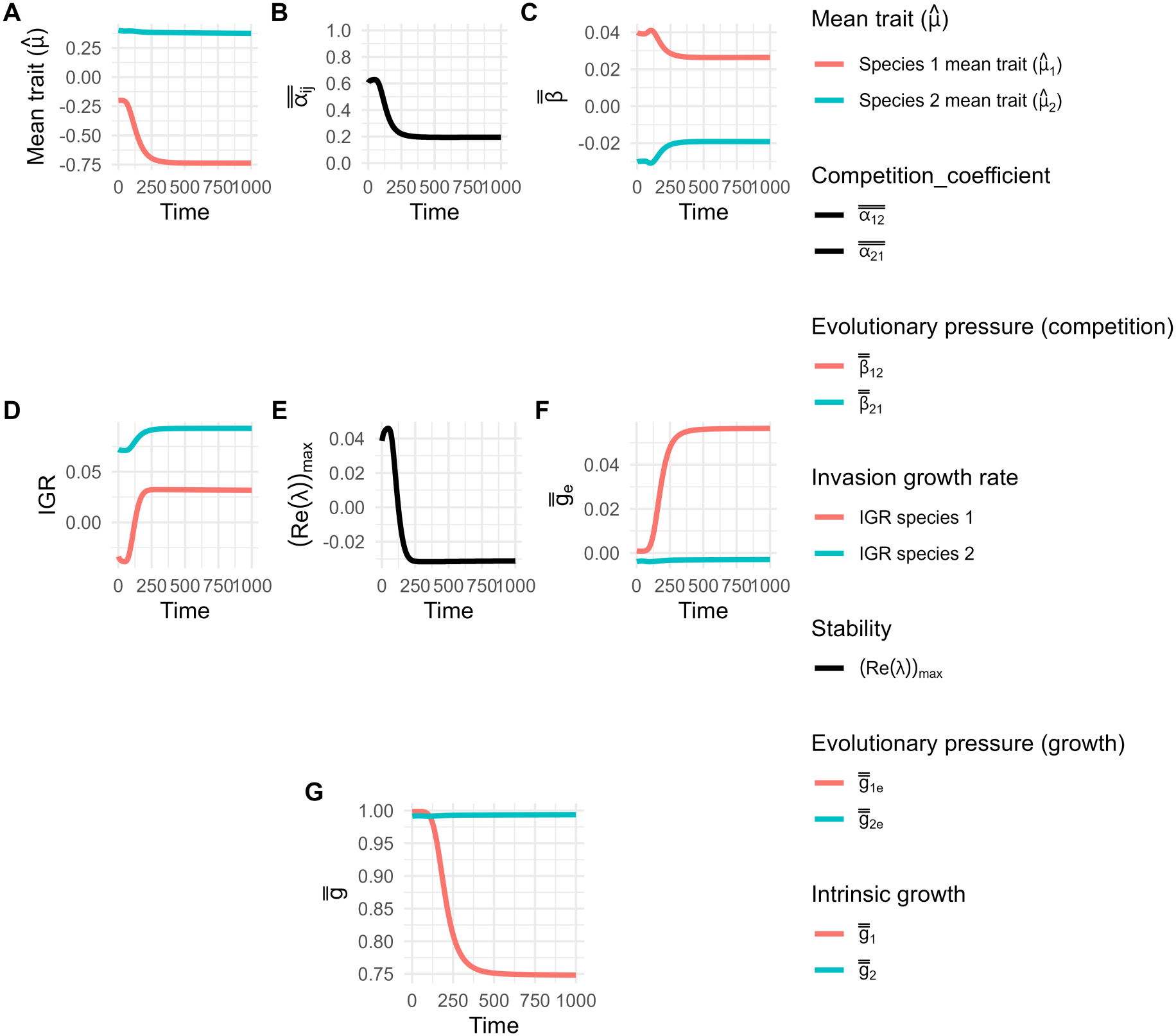
Detailed dynamics for figure 3C-Evolution with convergent plasticity in a narrow environment: (**A**) Dynamics of mean traits. (**B**) Dynamics of competition coefficient (= niche overlap). (**C**) Dynamics of evolutionary pressure due to competition. (**D**) Dynamics of invasion growth rate. (**E**) Dynamics of the dominant eigenvalue of the community matrix (a negative value denotes stability equivalent to coexistence for a 2 species system; see supplementary section S1.5).(**F**) Dynamics of the evolutionary pressure due to growth. (**G**) Intrinsic growth rate. The time *t* (abscissa) represents the generations.

**Figure S14:**
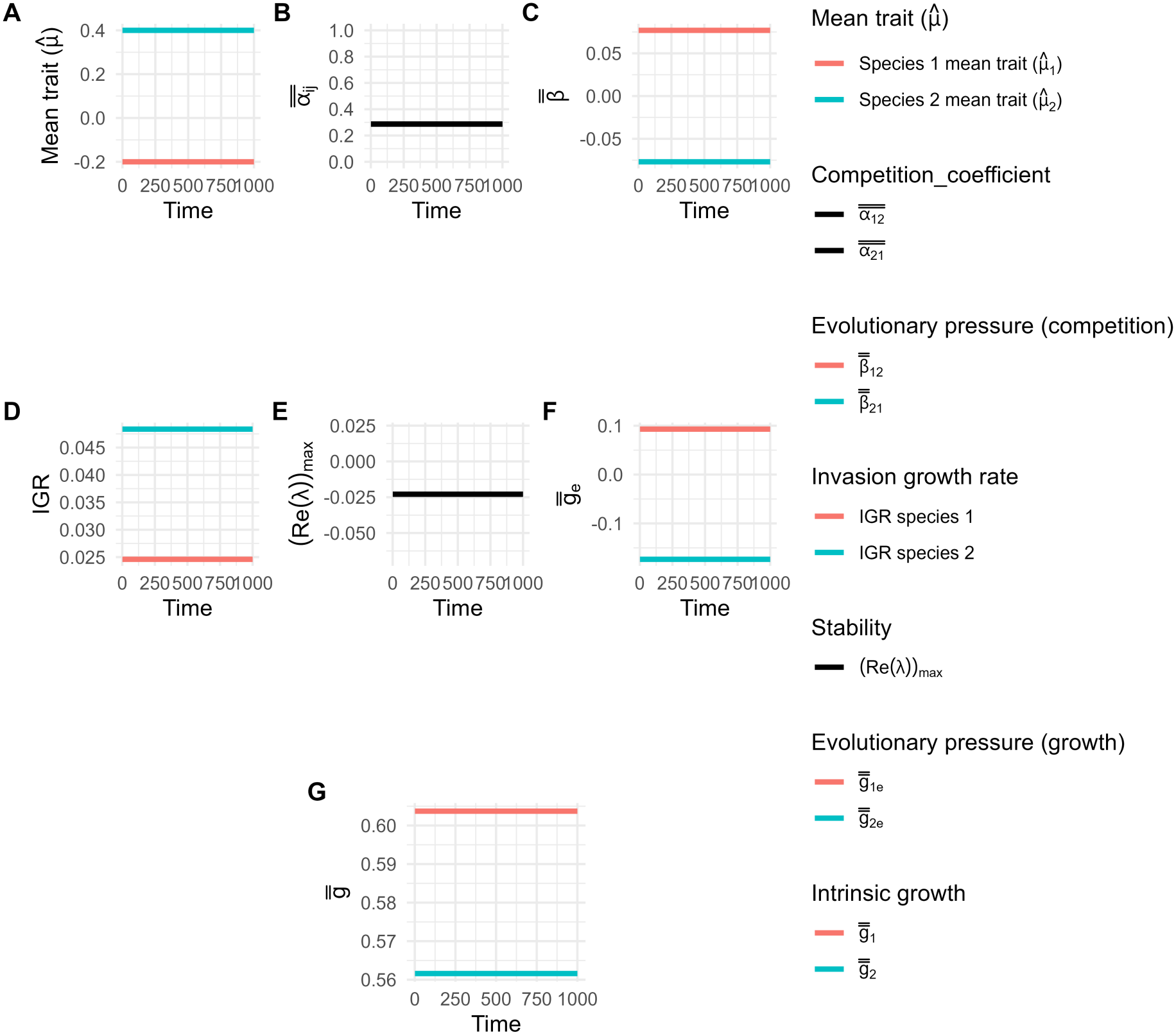
Detailed dynamics for figure 3D with no plasticity and no evolution with high ITV in a narrow environment: (**A**) Dynamics of mean traits. (**B**) Dynamics of competition coefficient (= niche overlap). (**C**) Dynamics of evolutionary pressure due to competition. (**D**) Dynamics of invasion growth rate. (**E**) Dynamics of the dominant eigenvalue of the community matrix (a positive value denotes instability equivalent to no coexistence for a 2 species system; see supplementary section S1.5). (**F**) Dynamics of the evolutionary pressure due to growth. (**G**) Intrinsic growth rate. The time *t* (abscissa) represents the generations.

**Figure S15:**
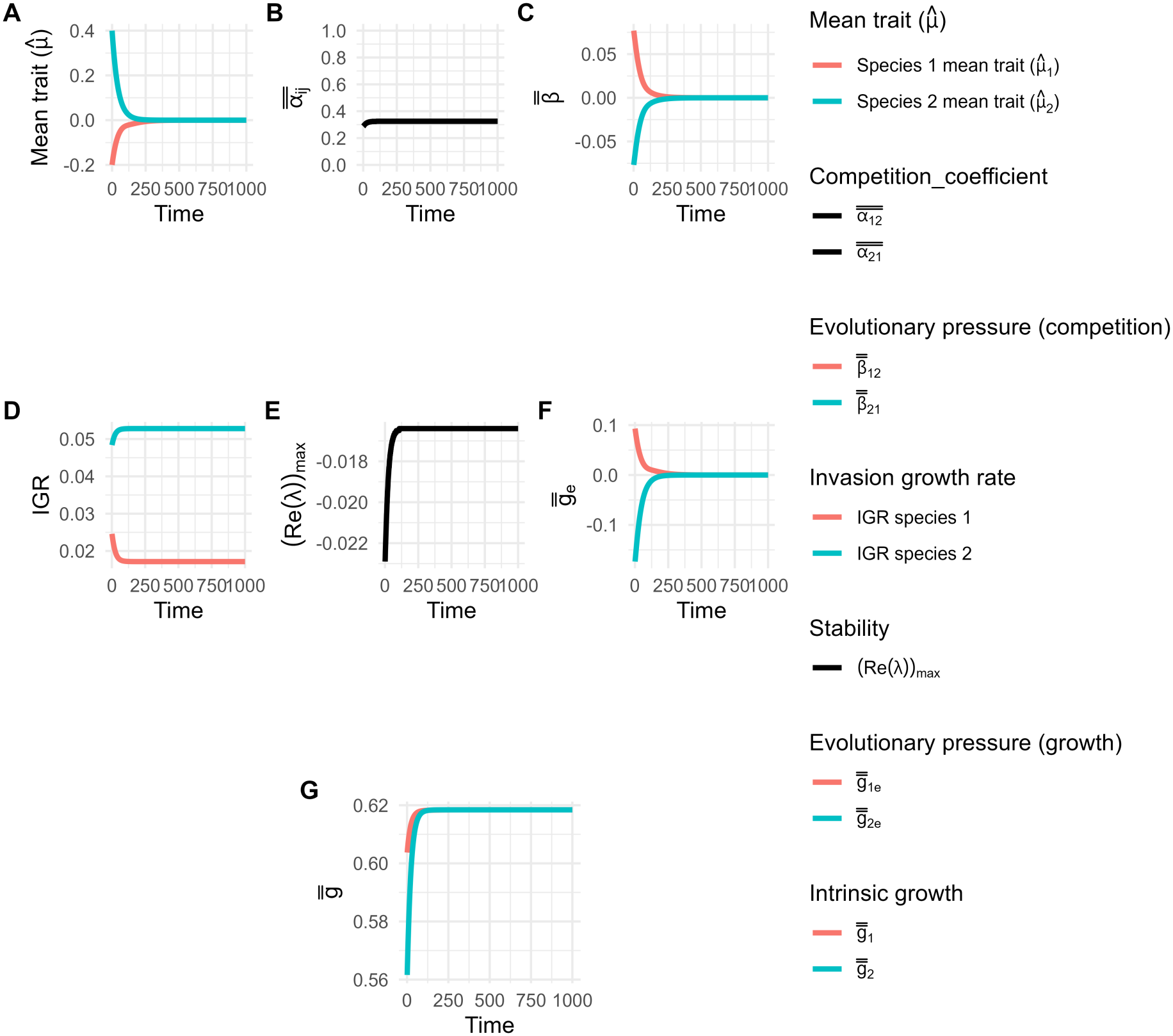
Detailed dynamics for figure 3E with only evolution with high ITV in a narrow environment: (**A**) Dynamics of mean traits. (**B**) Dynamics of competition coefficient (=niche overlap). (**C**) Dynamics of evolutionary pressure due to competition. (**D**) Dynamics of invasion growth rate. (**E**) Dynamics of the dominant eigenvalue of the community matrix (positive value denotes instability equivalent to no coexistence for a 2 species system; see supplementary section S1.5). (**F**) Dynamics of the evolutionary pressure due to growth. (**G**) Intrinsic growth rate. The time *t* (abscissa) represents the generations.

**Figure S16:**
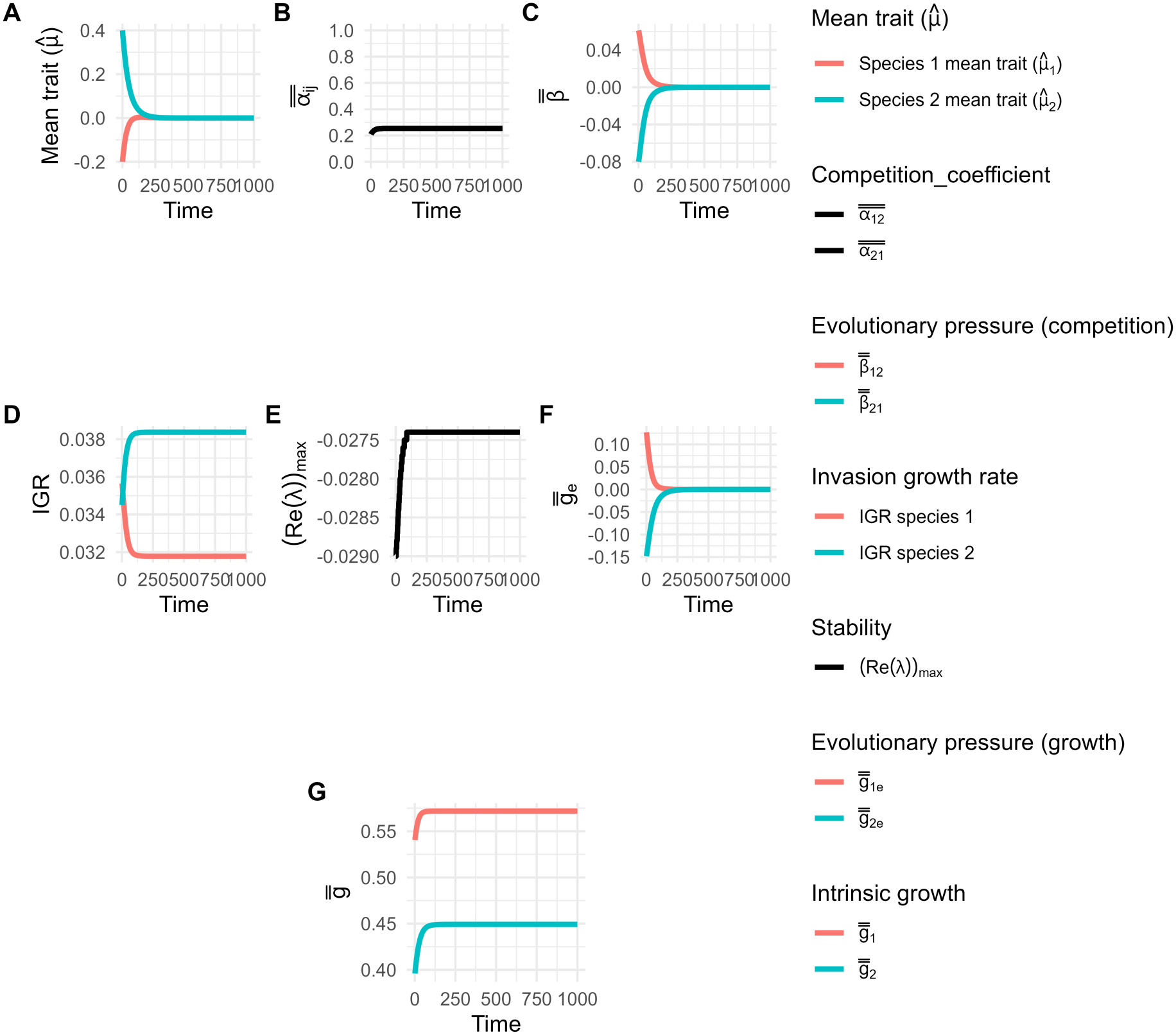
Detailed dynamics for figure 3F (green lines) with divergent plasticity and evolution with high ITV in a narrow environment: (**A**) Dynamics of mean traits. (**B**) Dynamics of competition coefficient (= niche overlap). (**C**) Dynamics of evolutionary pressure due to competition. (**D**) Dynamics of invasion growth rate. (**E**) Dynamics of the dominant eigenvalue of the community matrix (a positive value denotes instability equivalent to no coexistence for a 2 species system; see supplementary section S1.5). (**F**) Dynamics of the evolutionary pressure due to growth. (**G**) Intrinsic growth rate. The time *t* (abscissa) represents the generations.

**Figure S17:**
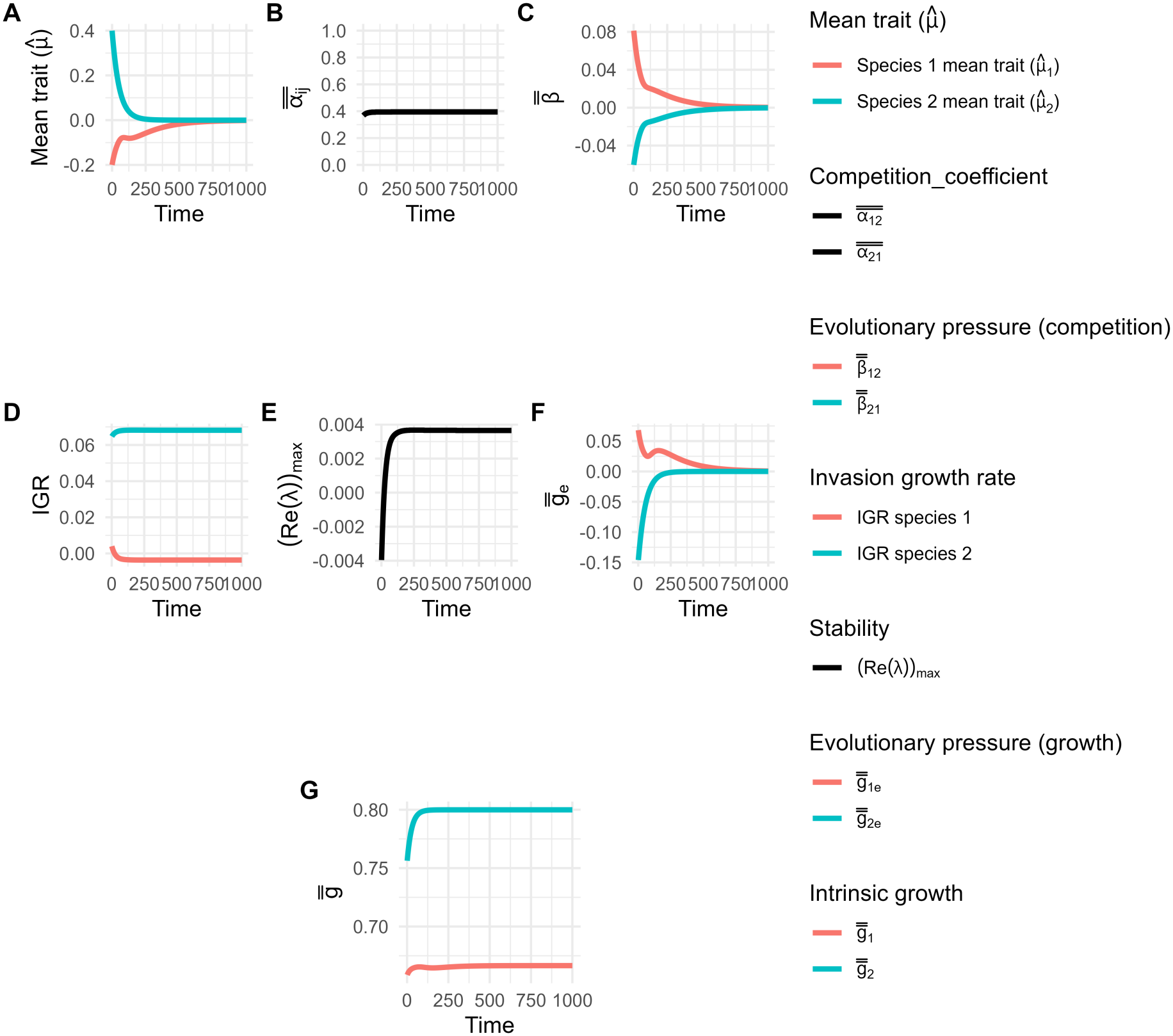
Detailed dynamics for figure 3F (pink lines) with convergent plasticity and evolution with high ITV in a narrow environment: (**A**) Dynamics of mean traits. (**B**) Dynamics of competition coefficient (= niche overlap). (**C**) Dynamics of evolutionary pressure due to competition. (**D**) Dynamics of invasion growth rate. (**E**) Dynamics of the dominant eigenvalue of the community matrix (positive value denotes instability equivalent to no coexistence for a 2 species system; see supplementary section S1.5). (**F**) Dynamics of the evolutionary pressure due to growth. (**G**) Intrinsic growth rate. The time *t* (abscissa) represents the generations.

**Figure S18:**
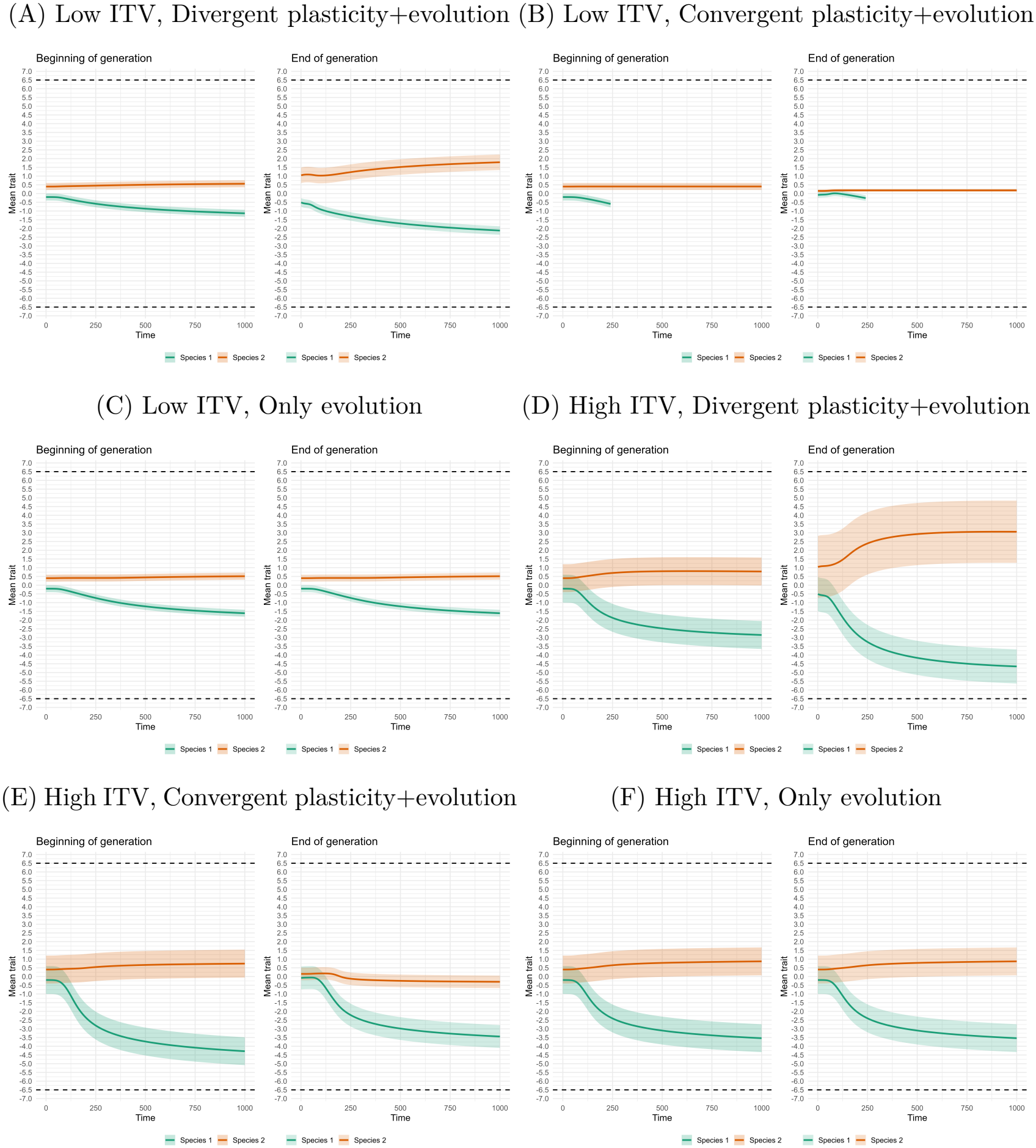
Mean trait dynamics for broad environments (figure 2 scenarios, *θ* = 6.5, *ω* = 1.1): Dynamics of the mean trait of each species for various scenarios, as given by (**A**)-(**F**). In each subplot, the first panel denotes the mean trait value at the beginning of the generation and the second panel denotes the mean trait value at the end of that respective generation after the plastic shift of traits. The dashed lines denote the abiotic environment breadth. The error bars represent the species level trait standard deviation *σ*.

**Figure S19:**
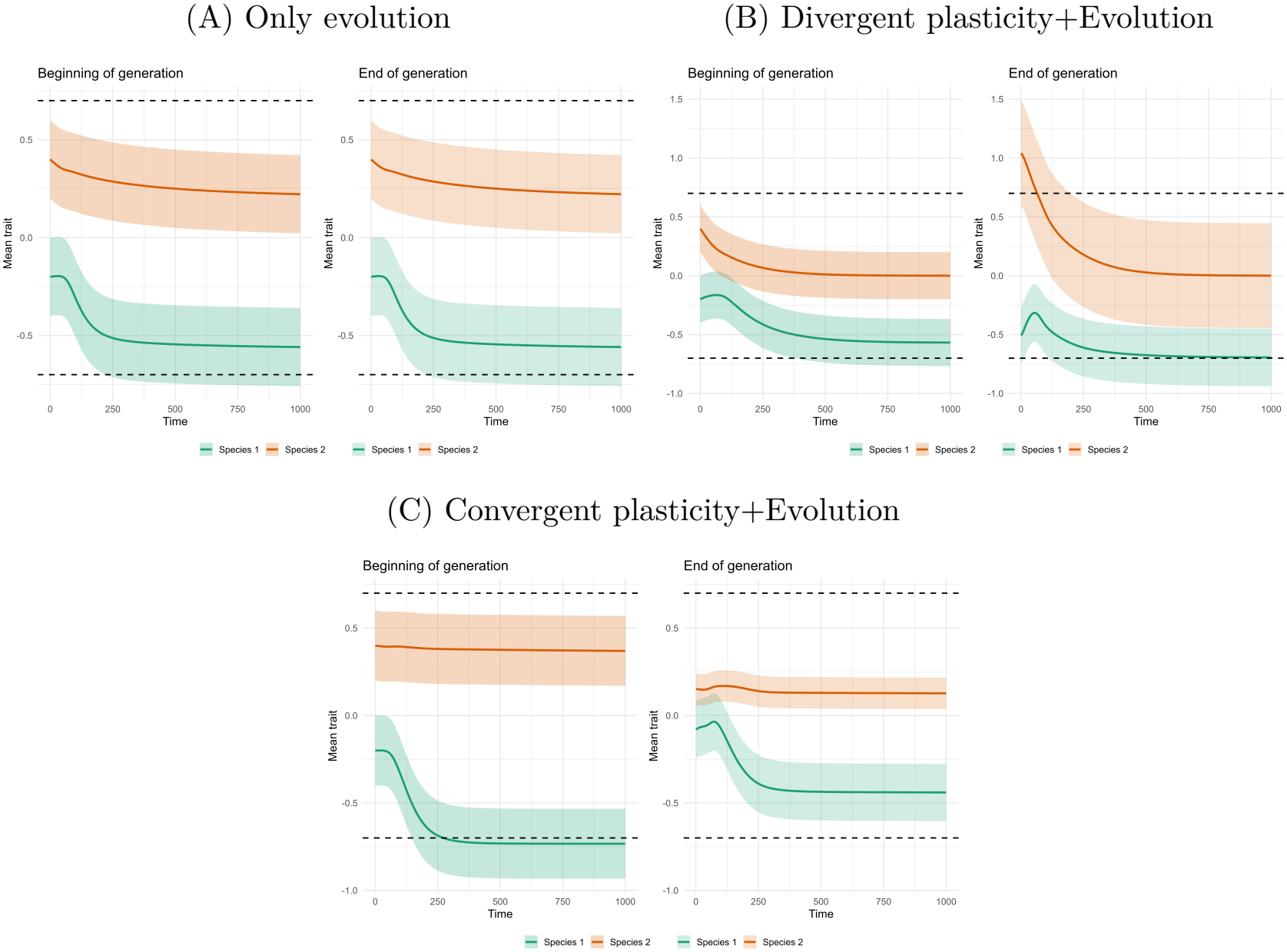
Mean trait dynamics for narrow environments and low ITV scenarios (figure 3A-C scenarios, *θ* = 0.7, *ω* = 0.55): Dynamics of the mean trait of each species for various scenarios, as given by (**A**)-(**C**). In each subplot, the first panel denotes the mean trait value at the beginning of the generation and the second panel denotes the mean trait value at the end of that respective generation after the plastic shift of traits. The dashed lines denote the abiotic environment breadth. The error bars represent the species level trait standard deviation *σ*.

**Figure S20:**
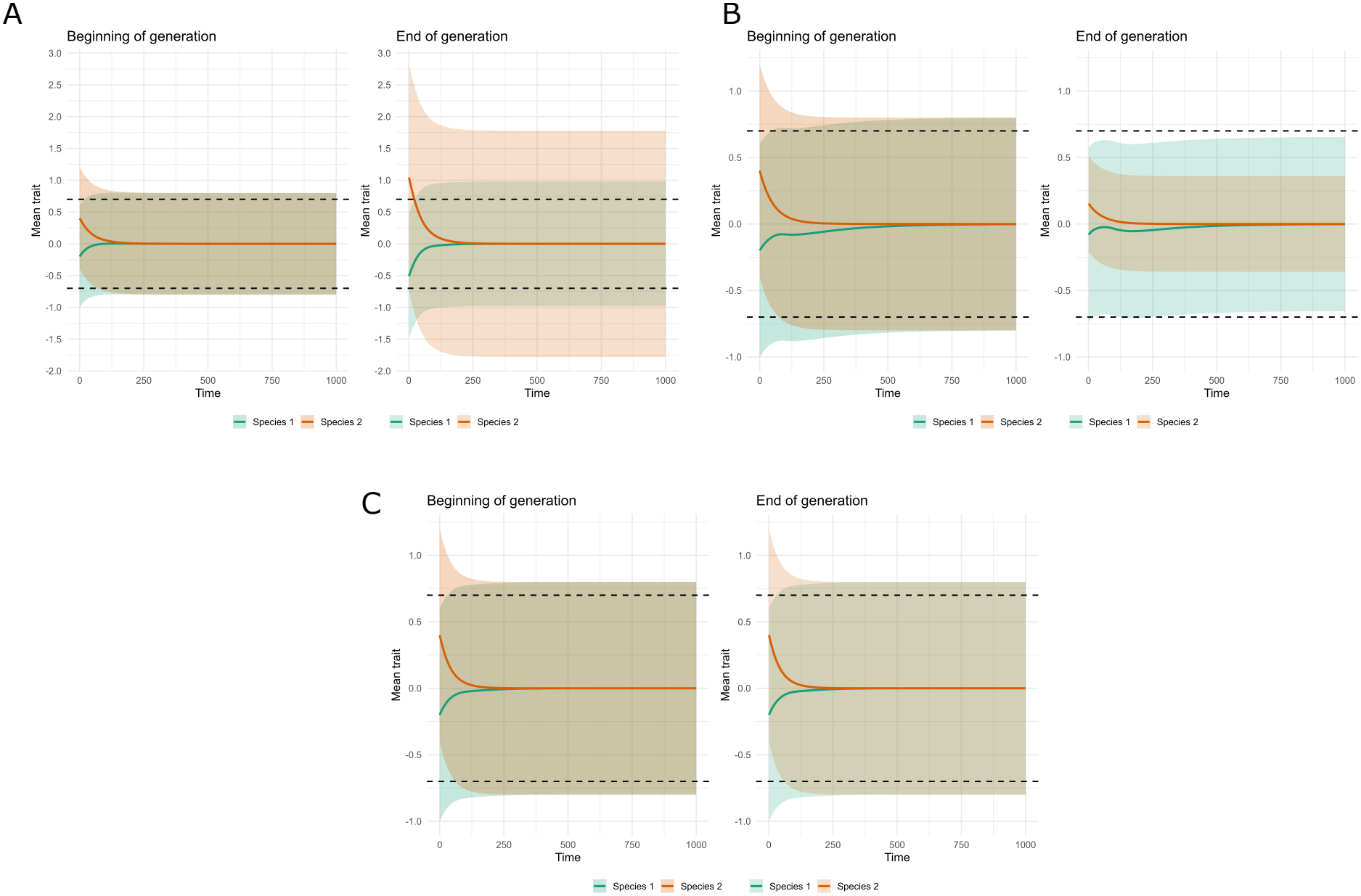
Mean trait dynamics for narrow environments and high ITV scenarios (figure 3D-F scenarios, *θ* = 0.7, *ω* = 0.55): Dynamics of the mean trait of each species for various scenarios: (**A**) divergent plasticity and evolution, (**B**) convergent plasticity and evolution and (**C**) evolution without plasticity. In each subplot, the first panel denotes the mean trait value at the beginning of the generation and the second panel denotes the mean trait value at the end of that respective generation after the plastic shift of traits. The dashed lines denote the abiotic environment breadth. The error bars represent the species level trait standard deviation *σ*.

**Figure S21:**
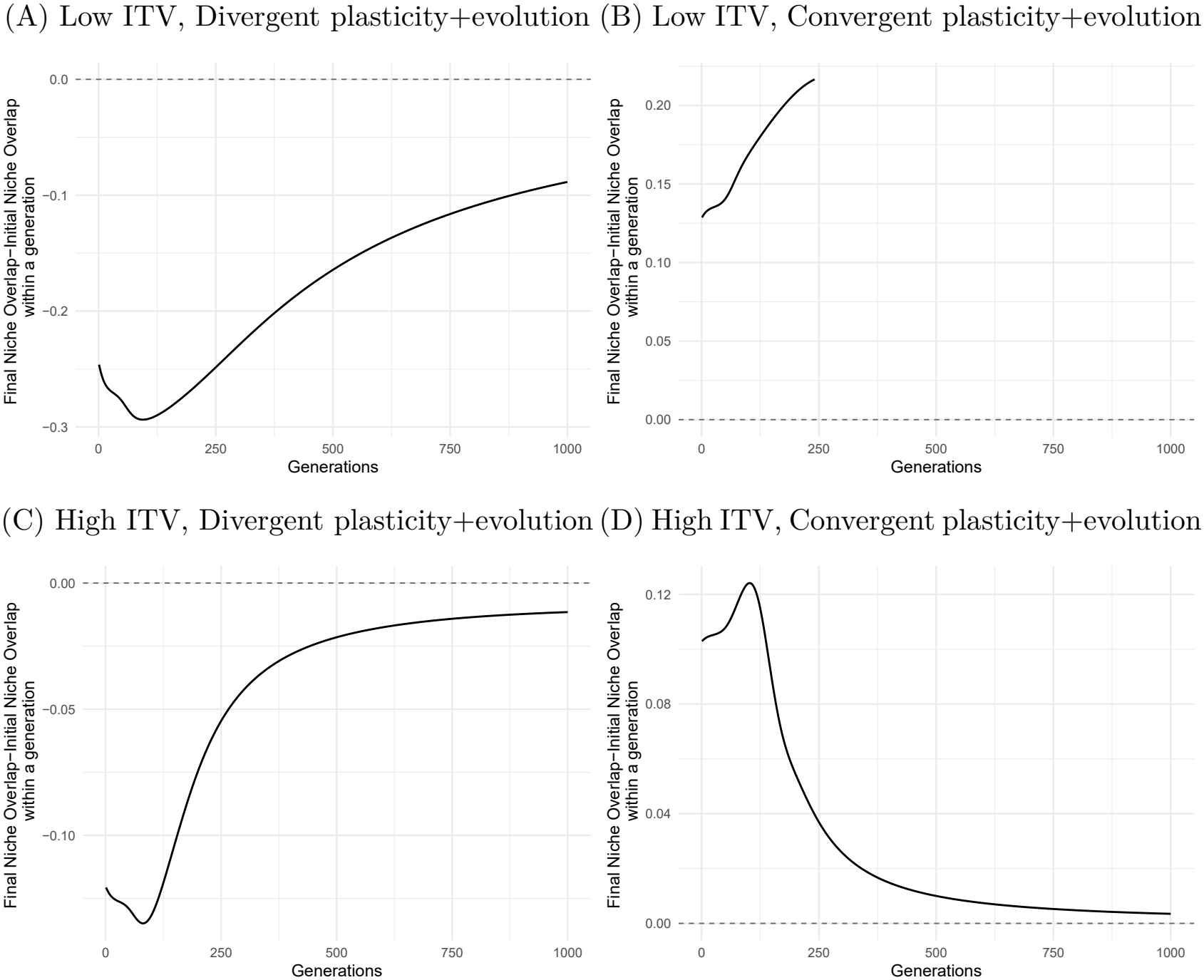
Within-generation effect of plasticity across generations in broad environment (figure 2 scenarios, *θ* = 6.5, *ω* = 1.1): The difference between the final niche overlap (after the plastic trait shift) and the initial niche overlap. Positive values denote within-generation trait distribution convergence (convergent plasticity), and negative values denote within-generation trait distribution divergence (divergent plasticity).

**Figure S22:**
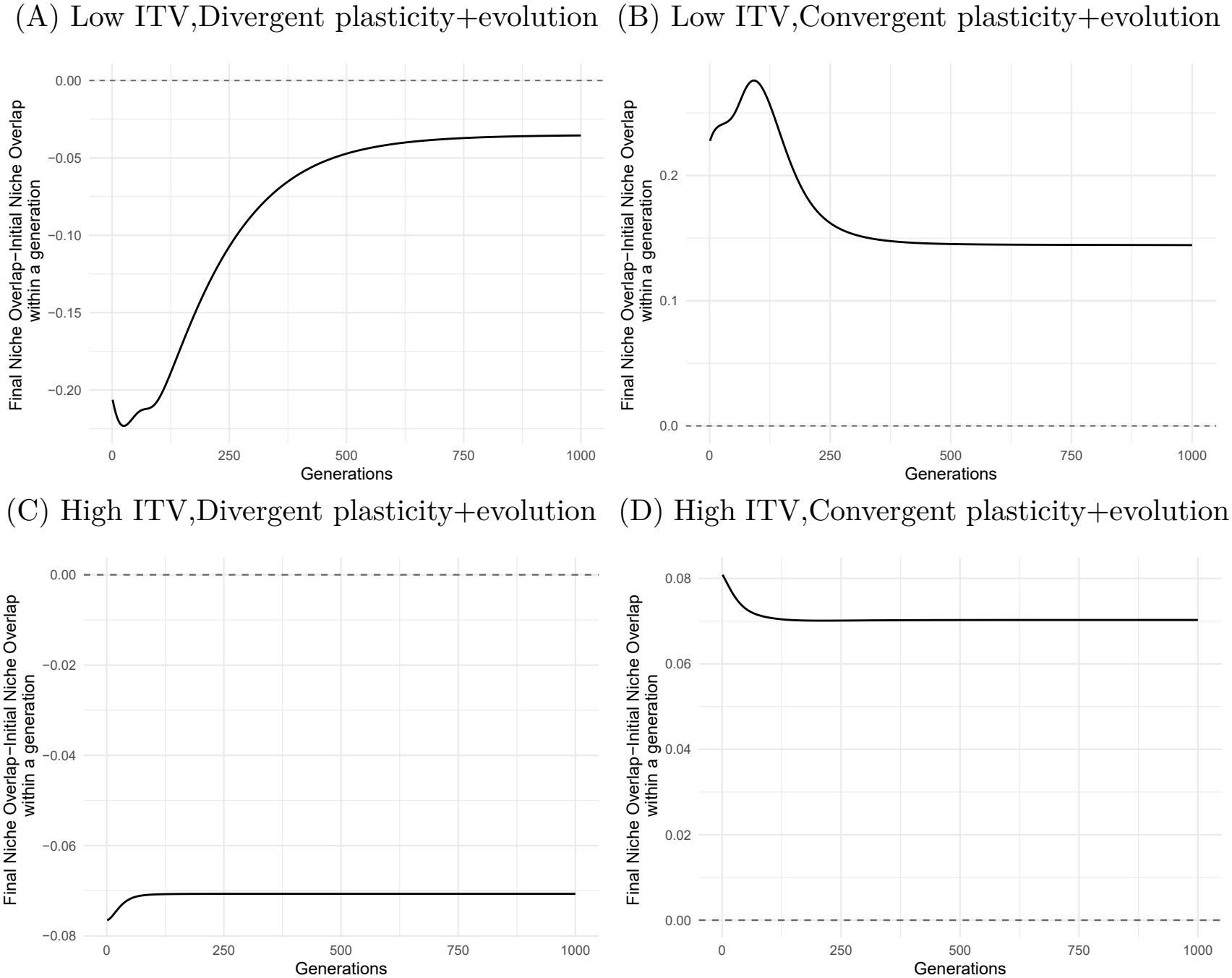
Within-generation effect of plasticity across generations in a narrow environment (figure 3 scenarios, *θ* = 0.7, *ω* = 0.55): The difference between the final niche overlap (after the plastic trait shift) and the initial niche overlap. Positive values denote within-generation trait distribution convergence (convergent plasticity), and negative values denote within-generation trait distribution divergence (divergent plasticity).

### S1.7 Effect of plasticity on the speed of evolution

The debate of whether phenotypic plasticity facilitates or hinders adaptive evolution has largely focused on adaptation to abiotic environments. Plasticity is thought to slow evolution when the plastically modified trait is closer to the optimum trait, thereby reducing the evolutionary pressure (Ghalambor et al., 2007). Plasticity can speed up evolution when the plastically modified trait is away from the optimum trait, thereby increasing the evolutionary pressure (Ghalambor et al., 2015). Plasticity can facilitate evolution in two ways. First, by increasing persistence under stressful conditions, it can prevent extinction and “buy time” for genetic adaptation. This is termed as buying time hypothesis (Fox et al., 2019; Júnior et al., 2025). Second, according to the plasticity-first hypothesis, the plastic responses can generate novel phenotypes on which selection can act. Selection favors individuals whose expressed phenotype is closer to the optimum (Levis & Pfennig, 2016).

In the case of two competing species (in a constant abiotic environment) as in our model, the above-mentioned general analysis might not readily apply as each species is the biotic environment for the other species and thus the species reciprocally act as the selective environment for each other. For example, in the case of a broad environment (figure S23) (where evolution tries to reduce only competition), we find that the effect of plasticity on the speed of evolution of each species can be asymmetric, i.e., the same plastic response can increase the rate of evolution in one species and decrease the rate of evolution in the competing species. By speed of evolution, we mean, 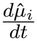. In this, 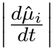 represents the magnitude of speed of evolution, while the sign indicates the direction in which the mean trait is evolving ( a positive sign indicates that the trait evolves towards +*∞* and a negative sign indicates that it evolves towards *−∞*). For example, if 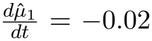 when 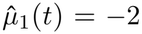, and 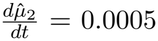 when 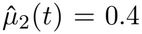, then the magnitude of evolutionary change is greater for species 1 than for species 2. Thus, species 1 evolves rapidly to more negative trait values, whereas species 2 evolves slowly to more positive trait values.

To better understand this phenomenon, we now present a detailed analysis of a scenario for divergent plasticity and evolution in broader environment for a case of high ITV. For the weaker species 1, the speed of evolution was least with divergent plasticity and highest with convergent plasticity, whereas for the dominant species 2 it was the opposite (figure S23G-H). This is because, from Eq. 9, the speed of evolution of a species is dependent on the strength of selection (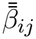, evolutionary pressure on species *i* due to species *j*) and the density of the competing species (*N*_*j*_), that is 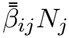. With divergent plasticity, the competition coefficient reduces (for both species). This allows the density of the weaker species to increase (figure S23C). Plasticity changes the trait distribution of each species within a generation (as evident from Eq. S47 and Eq. S48). Due to this, compared to the case of no plasticity (purple lines in figure S23A-B), with divergent plasticity (green lines), the evolutionary pressure on the weaker species (species 1) reduces (absolute values of 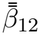 in figure S23A), but that on the dominant species (species 2) increases (absolute values of 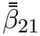 in figure S23B). This is because divergent plasticity (plasticity in general) affects asymmetry in the competitive effects experienced by individuals with different trait values within each species, reducing this asymmetry for the weaker species but increasing it for the dominant species (i.e., due the change in shape of the trait distribution of each species due to plasticity, for the weaker species, competition declines less as trait values move away from the competitor, whereas for the dominant species it declines more strongly; This is captured by the covariance structure of 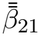 and 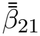 in Eq. S70 and Eq.S71). Overall, the increased magnitude of 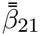 and the increased density of the weaker competitor, *N*_1_, increase the magnitude of selection term 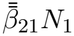 (figure S23F). Thus, compared to the case of only evolution, with divergent plasticity and evolution, the dominant species 2 evolves greater trait divergence (figure S23J). On the other hand, for the weaker species (species 1), compared to the case of only evolution, with evolution under divergent plasticity, the magnitude of 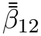 decreases (figure S23A), while *N*_2_ changes negligibly (figure S23D). Consequently, the magnitude of the selection term, 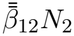, decreases, resulting in a lower speed of evolution compared with evolution alone (figure S23E).

The magnitude of total evolutionary change of the mean of each species is dictated by the push of the competing species (from Eq. 9, with the selection term maximizing intrinsic growth rate being zero for large environmental breadth). Since the density of the dominant species (*N*_2_) is very high compared to the weaker species *N*_1_, the weaker species’ trait mean evolves more than the dominant species’ trait mean (figure S23I-J).

**Figure S23:**
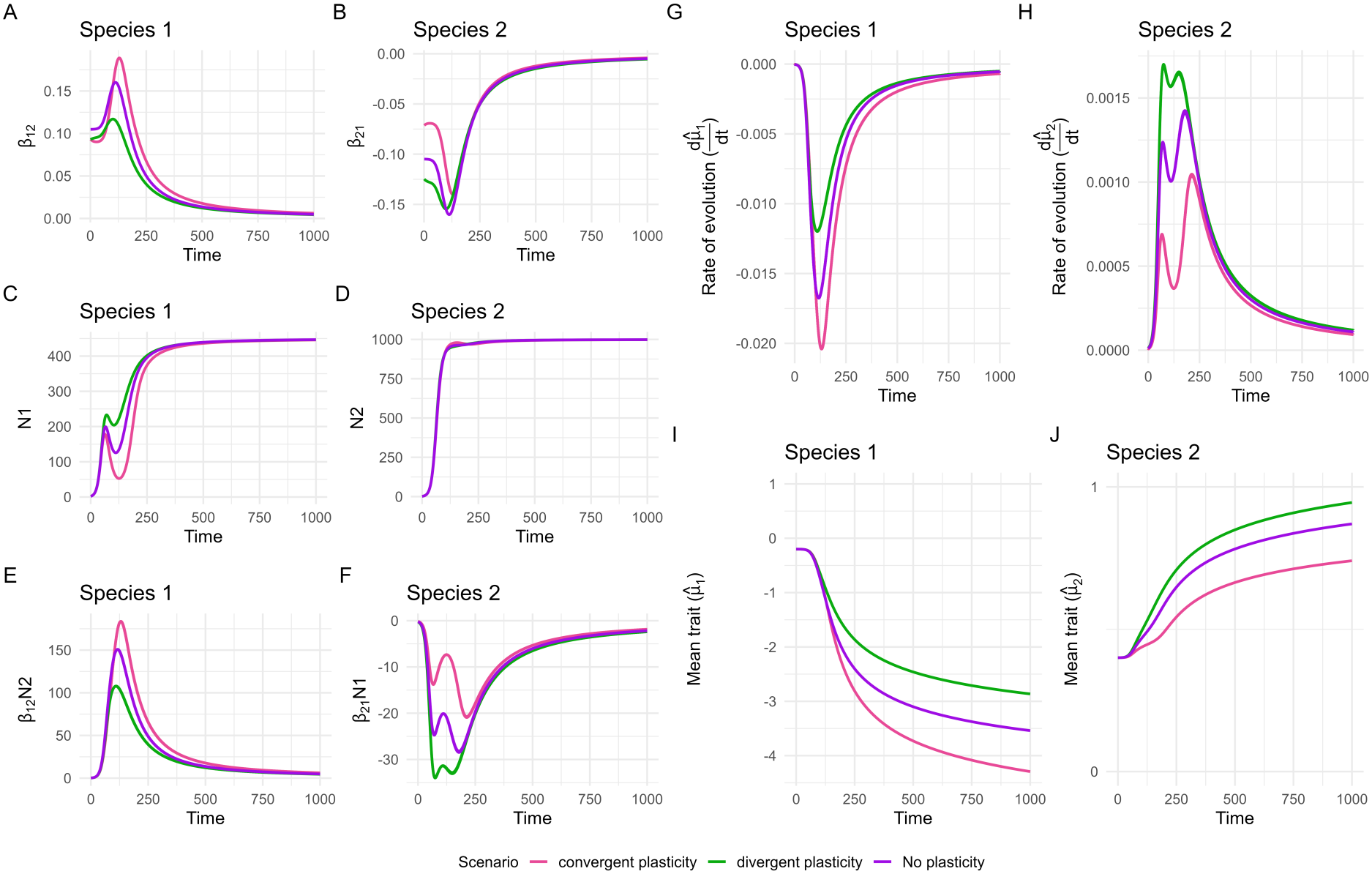
Effect of phenotypic plasticity on the rate of evolution in a broad environment. (**A**)-(**F**): Constituents of the rate of evolution (at the species level) as per Eq. 9 for various plasticity scenarios. (**G**)-(**H**): Dynamics of rate of evolution (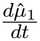 and 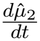) for various plasticity scenarios. (**I**)-(**J**): Dynamics of mean traits of each species. Parameter values used: *K*_1_=450, *K*_2_=1000, 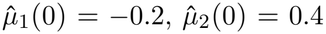, *σ*_1_ = *σ*_2_ = 0.8, *θ* = 15, *ω* = 1.1. Due to very high environmental breadth, 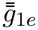 and 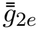 are 0 and thus do not affect evolution.

### S1.8 Adaptive and maladaptive nature of plasticity

The effect of plasticity is considered adaptive if the direction of change in trait due to plasticity within a generation aligns with the direction due to evolution, and maladaptive if the direction is aligned opposite in that environment (Ghalambor et al., 2015). However, this criterion may not be readily transferable to eco-evolutionary competitive dynamics, as in our model, where the environment being biotic is not fixed.

In competing species, the selective environment for a species changes with competitors traits and abundances. In such a scenario, selection-driven evolutionary change of a species does not necessarily ensure population persistence under competitive and abiotic constraints and thus could also result in exclusion. This also means that a plastic response aligned to direction of such evolutionary response also cannot ensure population persistence. For example from our results, in narrow environments with low ITV, plastic and evolutionary trait changes occur in the same direction in the weaker species under divergent plasticity (figure S19B), yet this species is excluded. With high ITV, these changes occur in opposite directions in both species under divergent plasticity (figure S20A), yet their long-term abundances exceed those under evolution alone. These outcomes demonstrate that directional alignment does not necessarily predict persistence or abundance, and thus not establish the adaptive value of plasticity. Assessing whether plasticity is adaptive or maladaptive thus requires comparing fitness with and without the focal plastic response under the same abiotic and competitive conditions.

